# The UPR drives gametogenesis via an unexpected response, coordinating translation and ER structure

**DOI:** 10.64898/2026.05.29.728519

**Authors:** Constantine Bartolutti, Alena Bishop, Yanzhe Ma, Kelsey Van Dalfsen, Elçin Ünal, Marko Jovanovic, Gloria Ann Brar

## Abstract

Stress responses, including the unfolded protein response (UPR), are commonly studied via induction with harsh exogenous stressors, leaving endogenous functions of these pathways less well understood. We found that the endogenous UPR that precedes meiosis in budding yeast is required for gamete production but diverges dramatically from previously defined UPR outputs, with only a few characterized UPR targets induced, and mildly. The role of this UPR can be replaced by increasing ER chaperones, reducing bulk translation, or impairing the machinery for protein translocation into the ER. ER integrity appears compromised in pre-meiotic cells lacking the UPR, as foci of reticulon proteins are seen and correlate strongly with an inability of cells to enter meiosis. These findings indicate that physiological UPR activation supports proteostasis and normal ER structure, preparing cells for meiotic entry by reducing the load of proteins that enter the ER. Overall, our study reveals surprising features of a physiological UPR induction that enables a cell fate decision.

## Introduction

Approximately one-third of proteins produced by cells are translocated into the endoplasmic reticulum (ER), a proportion that varies with cell type and condition, necessitating a mechanism for cells to safeguard and modulate their ER protein folding capacity (Brodsky and Skach, 2011; Aviram and Schuldiner, 2017). The Unfolded Protein Response (UPR) is a conserved pathway that mediates this, resulting in upregulation of pro-folding factors, including chaperones (most famously BiP/GRP78, Kar2 in yeast), quality control factors, and redox enzymes (Walter and Ron, 2011). The UPR was first discovered in budding yeast as a signaling pathway that could be stimulated in response to external triggers, including the drug dithiothreitol (DTT), leading to bulk misfolding of ER-resident proteins. Accumulation of misfolded proteins leads to activation of the ER membrane protein Ire1, which transduces the signal by mediating cytoplasmic splicing of the mRNA for a single target, XBP-1 in metazoans and *HAC1* in yeast (Sidrauski et al., 1996; Chapman et al., 1998). The protein produced from this spliced mRNA is a transcription factor that has been shown to drive transcription of hundreds of target mRNAs, including BiP/Kar2 (Cox et al., 1993; Cox and Walter, 1996; Mori et al., 1998, 1996; Travers et al., 2000). Beyond this shared signaling from yeast to human, there are additional features of the UPR in higher eukaryotes that are absent in budding yeast. These include the ability of Ire1 to drive degradation of mRNAs through RIDD (regulated Ire1-dependent decay) and the presence of two additional UPR branches, mediated through the transcription factor ATF6 or the kinase PERK (Hollien and Weissman, 2006; Hollien et al., 2009; Maurel et al., 2014; Walter and Ron, 2011). These additional UPR functionalities drive, among other things, downregulation of bulk protein synthesis, which has not been previously reported for the IRE-1 branch of the UPR (Walter and Ron, 2011).

Most studies of the UPR have focused on the events that occur following experimental application of acute exogenous stressors that stimulate ER protein misfolding. The UPR is also known to be important in physiological states in metazoans, including for the differentiation of antibody-producing cells and embryonic development, but much less is known about how the UPR is activated under such natural conditions and the specific physiological output that it drives (Reimold et al., 2001, 2000). We recently found that the UPR is important for basal growth of wild-type (WT) budding yeast cells (Bartolutti et al., 2025). We found that the role of the UPR in maintaining normal ER function in unstressed conditions is important enough that adaptive aneuploidy invariably occurs in cells lacking UPR activity, even if they are never exposed to exogenous stressors. The specific aneuploidies that occur upon UPR loss either increased expression of Kar2 or reduced expression of machinery involved in translocating proteins into the ER. Remarkably, these aneuploidies resulted in a profound growth rescue in UPR-deficient mitotic cells, despite also inducing cytosolic proteostatic stress. This study revealed a previously unrecognized physiological role for the UPR and demonstrated that maintaining ER protein folding capacity is critical for cellular fitness.

An additional physiological context in which we observed evidence for activation of the UPR was meiosis, the process by which gametes are produced (Brar et al., 2012). Meiosis has been traditionally defined by chromosomal events, including one round of DNA replication followed by two rounds of chromosome segregation, which together allow a diploid precursor cell to produce haploid gametes. It has recently become more widely recognized that wholesale remodeling and segregation of nearly every other cellular component is also crucial for meiotic success, and that this occurs in a manner that is coordinated with chromosome segregation (Suda et al., 2007; Hsu et al., 2017; King et al., 2019; Sawyer et al., 2019; Otto et al., 2021; Sing et al.; 2022, Suda et al., 2024; Xiao and Ünal, 2025). Here, we investigated the function of the UPR during pre- and mid-meiosis in budding yeast. We found that pre-meiotic UPR induction is critical for entry of cells into the meiotic differentiation program, and that its output diverges profoundly from the output defined in response to exogenous stressors. This endogenous UPR activation drives a very small number of transcriptional targets and appears to function primarily in supporting ER protein folding through downregulation of translation. Comparing the outputs of canonical UPR signaling to the pre-meiotic and mid-meiotic UPR output, we observe three fundamentally different effects on transcription, ranging from a broad transcriptional program focused on upregulating ER components to a distinct transcriptional response that results in reduced levels of ribosomal proteins, to a minimal response that is difficult to distinguish from uninduced cells. Our data suggest that the output of physiological pre-meiotic UPR is non-canonical relative to the established UPR program, and is critical for preparing cells to enter the meiotic program via support for normal ER structure and function.

## Results

### The UPR is internally activated in two waves as cells prepare for and undergo meiosis

Based on ribosome profiling data for cells undergoing meiosis, we previously reported endogenously driven translation of Hac1, the transcription factor that mediates the conserved Ire1-branch of the UPR in yeast (Brar et al., 2012). This occurred in two waves, with Hac1 translation in pre-meiotic cells, followed by downregulation when cells were transferred to conditions that stimulated meiotic progression and reactivation in mid-meiosis (late into the first meiotic division). Late into the second meiotic division, Hac1 translation again abated. This pattern of Hac1 translation was mirrored by patterns of splicing of the *HAC1* mRNA, and also by measurements of Hac1 protein abundance, indicating that Hac1 is unstable during meiosis (Figure 1A, 1B), as has been reported in ER stress of vegetative cells (Chapman and Walter, 1997; Kawahara et al., 1998). Both pre- and mid-meiotic waves of UPR activation occur and abate in the absence of external triggers that are known to activate the UPR, thus representing physiological induction and silencing.

**Figure 1:**
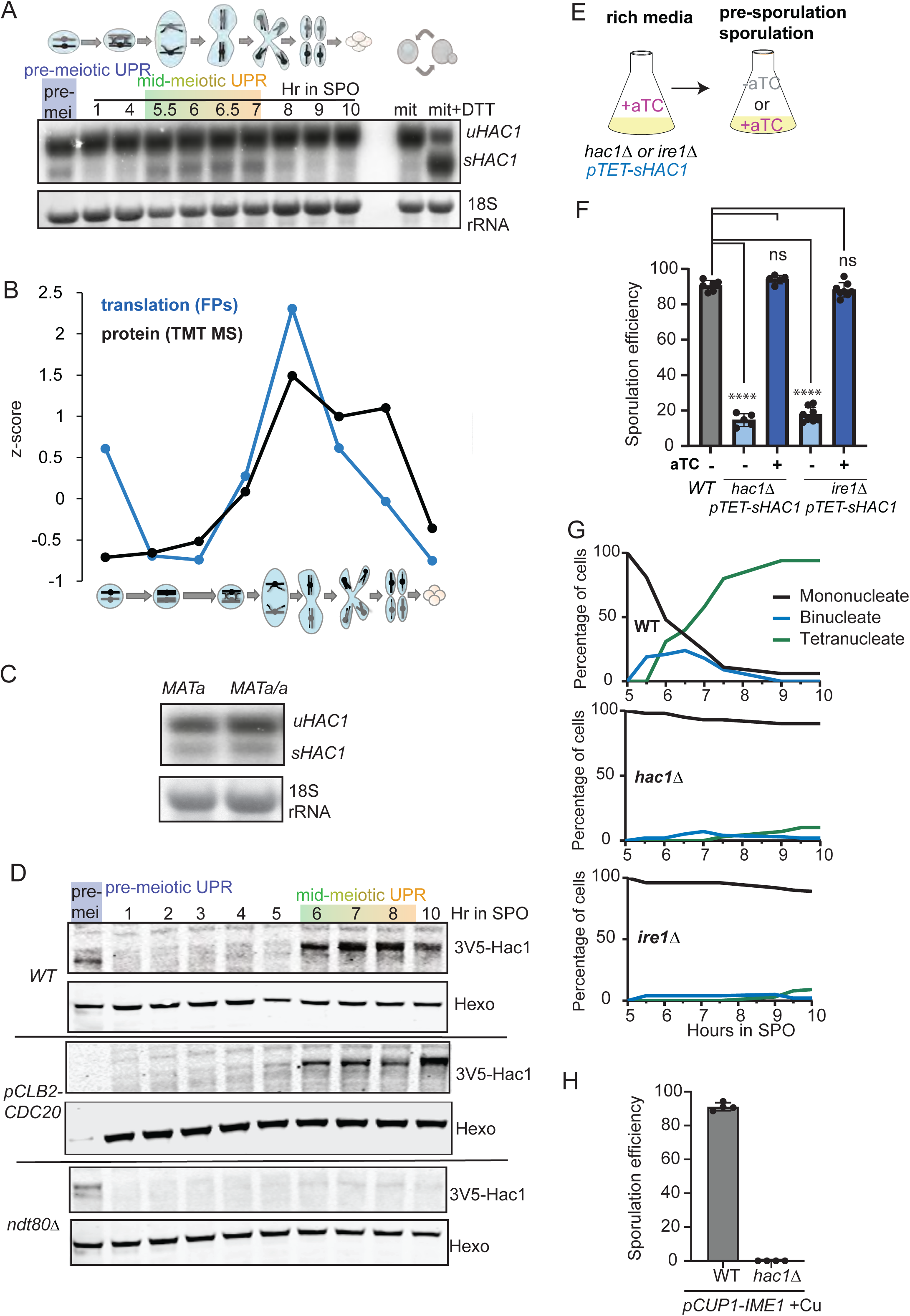
Physiological UPR induction is associated with and required for meiosis. (A) Northern blotting to detect unspliced (*u)* and spliced (*s) HAC1* mRNA in pre-meiotic and meiotic cells. Note the higher fraction of spliced *HAC1* observed with DTT treatment at right. (B) Ribosome profiling (blue) and TMT mass spectrometry (black) data for Hac1 from a previous study (Cheng et al., 2018) over timepoints during meiosis. (C) Northern blotting to detect unspliced (*u)* and spliced (*s) HAC1* mRNA in haploid cells (*MATa*) and diploid cells that are unable to enter meiosis (*MATa/MATa*) in pre-meiotic media. (D) Western blotting to assess Hac1 protein levels during a meiotic timecourse in WT, *pCLB2-CDC20*, and *ndt80Δ* cells. (E) Experimental scheme to maintain euploid UPR-deficient cells. Supplying them with aTC before meiosis drives *sHAC1* production. (F) Sporulation efficiency as scored by microscopy following 24 hours in sporulation media. 300 cells counted per strain/condition per experiment, minimum of n=5. (G) Meiotic progression scored by Mono-, bi-, and tetra-nucleate cells assessed by DAPI staining. 100 cells counted per timepoint and strain. (H) Sporulation efficiency as scored by microscopy following 24 hours in sporulation media with 50µM CuSO4 addition at 2h in SPO. 300 cells counted per strain/condition per experiment, n=4.

Efficient stimulation of meiosis in yeast requires pre-growth in media lacking fermentable carbon sources, and we observe *HAC1* splicing in pre-meiotic cells under these conditions (Figure 1A, 1B). To test whether this early wave of UPR induction is a result of early stages of meiosis or the media conditions used to stimulate them, we grew either haploid cells or diploid cells with two copies of the *MATa* locus and lacking the *MAT*α locus in pre-meiotic media. Neither of these cells is capable of entering meiosis, but we observed *HAC1* splicing in both cases, indicating that UPR induction under these conditions is stimulated by changes in growth media that are preparatory for meiosis, rather than a cue inherent to the early meiotic program (Figure 1C). During meiosis, diploid cells undergo homologous recombination, followed by two rounds of chromosome segregation, and generation of gametes (spores in budding yeast). A mutant that arrests at metaphase I and thus does not efficiently undergo the two divisions that typify meiosis shows a similar pattern of Hac1 protein abundance to WT cells, indicating that the normal meiotic chromosome segregation program is not needed to stimulate Hac1 production (Figure 1D; *pCLB2-CDC20;* Lee and Amon, 2003). We tested whether the mid-meiotic UPR induction is based on media conditions or the meiotic program by examining Hac1 protein levels in cells lacking the ability to exit meiotic prophase and undergo chromosome segregation (*ndt80Δ;* Xu et al., 1995). We found that in the absence of the gene expression program that allows mid-meiotic events including chromosome segregation, Hac1 is not expressed (Figure 1D). Together, these results indicate that the pre-meiotic wave of UPR activation is independent of meiosis per se, but the wave of UPR activation during mid-meiosis is controlled by the meiotic program of gene expression.

### Endogenous UPR activation is required for entry into meiosis

To assess the impact of UPR loss on gamete production, we used strains that lack either the endogenous *HAC1* or *IRE1* genes, and also house a genomically integrated cassette expressing a pre-spliced version of *HAC1* (*sHAC1*) that is under control of the *TET* promoter (*pTET;* Azizoglu et al., 2021; Bartolutti et al., 2025). When grown in the presence of anhydrotetracycline (aTC) or when aTC is withdrawn for up to 2 days, these cells remain euploid (Bartolutti et al., 2025). They are highly sensitive to DTT treatment in the absence of aTC, indicating full loss of UPR activity, but resistant in the presence of aTC, indicating a functional UPR (Figure S1A). If aTC is withdrawn for more than 2 days, the cell population becomes aneuploid, owing to the role for the UPR in maintaining basal growth and preventing aneuploidy in mitotically growing yeast cells (Bartolutti et al., 2025).

Cells homozygously deleted for either *HAC1* or *IRE1* and grown for 24-48 hours in the absence of aTC formed spores from only 10-20% of precursor cells, compared to roughly 90% in WT cells or controls grown in the presence of aTC (Figure 1E, 1F). This result indicated that UPR activation is functionally important for successful execution of the meiotic program. Analysis of the nuclear divisions by microscopy of cells carrying an Htb1-mCherry-encoding construct at sequential timepoints following transfer to sporulation media revealed that UPR-deficient cells were largely blocked prior to the first chromosome segregation stage (Figure 1G). RT-qPCR analysis of *IME1*, a transcript that is induced upon meiotic entry, indicated that cells lacking a functional UPR do not efficiently enter meiosis (Figure S1B). Forcing expression of *IME1* does not rescue the ability to enter meiosis in cells lacking Hac1, indicating that the crucial function of the UPR is not solely to induce this key factor (Figure 1H). In diploid cells lacking components of the ER-associated degradation (ERAD) pathway, spore formation proceeded normally, showing that the UPR’s requirement for meiotic entry is not related to its role in inducing ERAD (Figure S1C, S1D).

### Pre-meiotic ER chaperone expression can bypass the role of the UPR in meiosis

Diploid cells lacking the UPR are unable to enter meiosis (Figure 1F-H), thus precluding us from assessing the functional impact of mid-meiotic UPR activation. To overcome this, we added aTC to *hac1Δ* cells carrying a genomically integrated *pTET-sHAC1* construct during growth in pre-meiotic media, then washed out the aTC prior to transfer of cells to sporulation media, thus providing UPR activity only during pre-meiotic stages and not during mid-meiosis (Figure 2A, S1E, S1F). Surprisingly, we found that cells grown in this manner were able to sporulate with similar efficiency to WT cells, and that meiotic divisions occurred with similar timing to WT cells (Figure 2B, S1H). This indicated that UPR activation during mid-meiosis was dispensable, in contrast to its crucial role in pre-meiotic cells.

**Figure 2:**
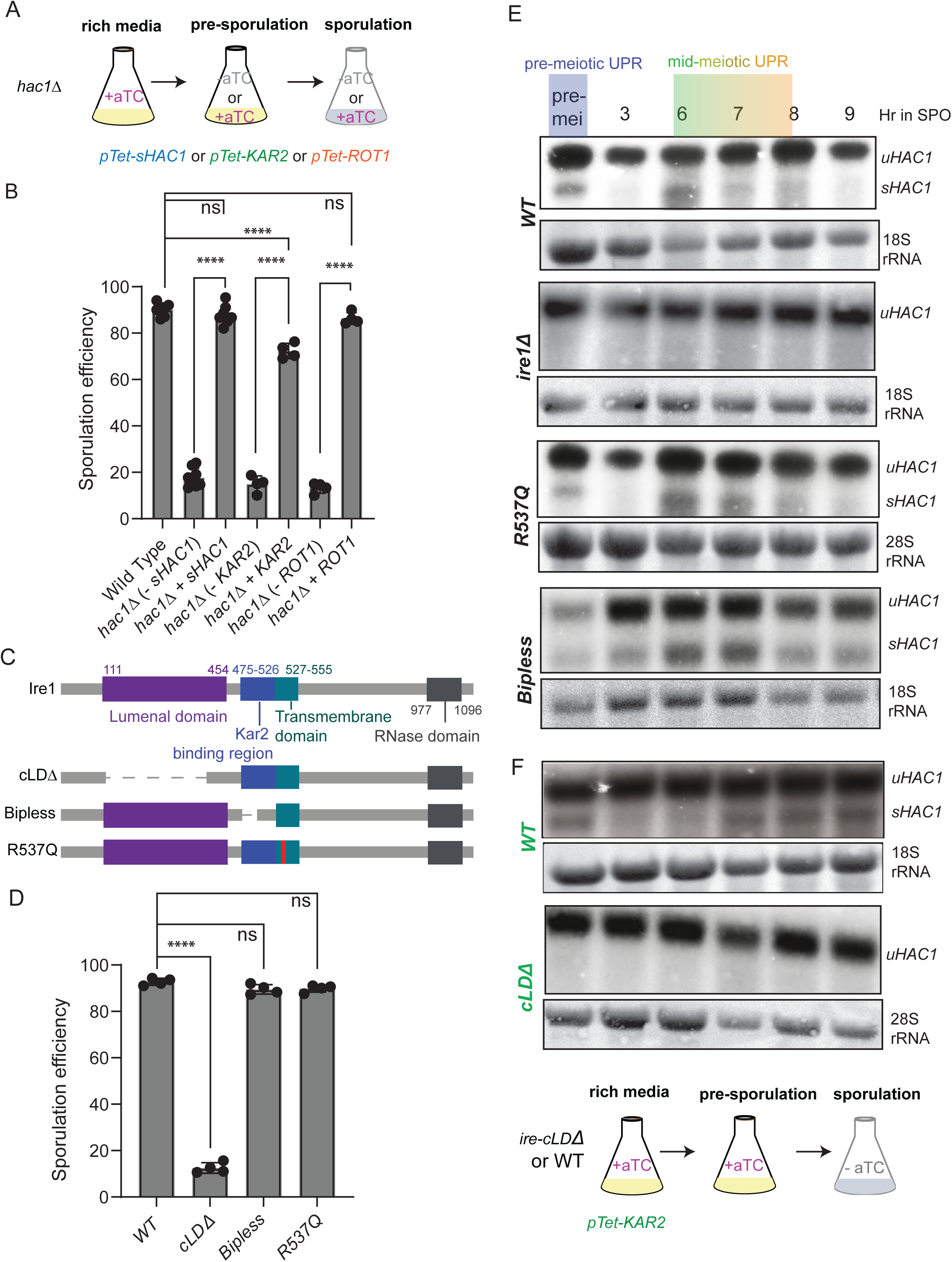
Demand for ER folding capacity is associated with pre-meiotic and mid-meiotic UPR activation. (A) Experimental strategy for supplying *hac1Δ* pre-meiotic cells with either *sHAC1, KAR2,* or *ROT1.* A concentration of 250ng/mL of aTC was used. (B) Sporulation efficiency as scored by microscopy following 24 hours in sporulation media following experimental strategy in (A). 300 cells counted per strain/condition per experiment, minimum of n=4. (C) Schematics of full-length Ire1 protein and mutants tested in (D) and (E). (D) Sporulation efficiency as scored by microscopy following 24 hours in sporulation media for cells in pre-sporulation and sporulation media without aTC. 300 cells counted per strain/condition per experiment, n=4. (E) Northern blotting to detect unspliced (*u)* and spliced (*s) HAC1* mRNA in pre-meiotic and meiotic cells. (F) *ire1-cLDΔ* containing cells required a different experimental strategy, based on overexpression *KAR2,* to enable mid-meiotic functionality assessments. Depicted below northern blots for this strain is a cartoon of that scheme, used also in a WT control. Significance was determined from unpaired t-test with Welch’s correction. P-value significance is **** < 0.0001, *** < 0.001, ** < 0.01., * < .05.

Our previous study of mitotic cells indicated that activation of only one UPR target, *KAR2*, was sufficient to compensate for loss of UPR activity (Bartolutti et al., 2025). We confirmed that additional Kar2 expression supported mitotic growth of UPR-deficient cells in the presence of DTT (Figure S1G). We found that providing *hac1Δ* cells with *KAR2* (via the same aTC-inducible system that we used with sHAC1; Figure S1A) was also able to substantially rescue spore formation, and that meiotic progression was normal in these cells (Figure 2B, S1H). We also tested whether Rot1, an essential ER protein which is not induced by the UPR, could rescue the ability to sporulate in cells lacking *HAC1*, using the same inducible system as we used for *KAR2* and *HAC1.*The specific role of Rot1 is unclear, with reports that it, like Kar2, acts as a general ER chaperone in *S. cerevisiae*, and that it has a role in glycosylation in *C. albicans*. (Figure 2B; Takeuchi et al., 2006; Takeuchi et al., 2008; Janik et al., 2026). Mutations in *ROT1* are synthetically lethal with those in *KAR2*, suggesting that it may support a similar role in supporting ER folding. Consistent with this model, its overexpression rescued sporulation in cells without Hac1, like the result for Kar2 (Figure 2B).

### Meiotic UPR induction depends on the lumenal peptide-sensing domain of Ire1

To determine what leads to UPR activation during mid-meiotic and pre-meiotic timepoints, we turned to analysis of Ire1. Ire1 has been shown to sense misfolded proteins within the ER lumen directly via a region in its lumenal domain, to sense lipid bilayer stress via its transmembrane domain (in particular residue R537), and to sense free lumenal levels of the ER chaperone Kar2 via a domain adjacent to the ER membrane (Credle et al., 2005; Promlek et al., 2011; Halbleib, 2017; Ho et al., 2020; Okamura et al., 2000). We constructed diploid cells with endogenous *IRE1* replaced with mutants to disrupt each of these sensing capabilities and assayed the ability of cells to complete meiosis and to internally activate splicing of *HAC1* (Figure 2C, 2D, 2E). We found that cells carrying the *ire1-R537Q* mutation, which are unable to sense lipid bilayer stress, displayed a WT pattern of *HAC1* splicing in pre-meiotic and mid-meiotic stages and completed meiosis with normal efficiency. Cells carrying the *ire1-bipless* allele (Pincus et al., 2010), in which Ire1 is unable to bind Kar2, completed meiosis normally and showed normal splicing of *HAC1,* but were unable to cease splicing after each wave in a timely manner (Figure 2D, 2F). This is consistent with the reported role for Kar2 binding to Ire1 serving a role to silence the activation of this sensor (Pincus et al., 2010). Cells carrying *ire1-cLDΔ* alleles (which are unable to bind misfolded peptides in the ER lumen) were unable to successfully complete meiosis (Figure 2D). To assess both pre- and mid-meiotic splicing of *HAC1* in these cells, we added a genomically inserted *pTET-KAR2* construct and grew cells in the presence of aTC until transfer to sporulation media. We found that these cells displayed no *HAC1* splicing at any pre-meiotic or meiotic stage despite completing meiosis with normal timing (Figure 2F). We concluded that pre-meiotic and mid-meiotic cells both activate the UPR in a manner that is dependent specifically on the ER lumenal peptide binding domain of Ire1. Together with the key role of ER chaperones in rescuing meiosis in UPR-deficient cells, these data indicate that the UPR acts prior to meiosis by bolstering the protein folding capacity within the ER lumen to prepare cells to enter meiosis.

### The mid-meiotic UPR does not induce a broad transcriptional program

In response to exogenous triggers, such as DTT or tunicamycin treatment, the UPR induces the transcription of hundreds of transcripts in concert in a Hac1-dependent manner (Travers et al., 2000). These transcriptional targets include those that encode ER chaperones (e.g. Kar2), ER redox factors (e.g. Pdi1 and Ero1), ERAD factors, and Hac1 itself, among many others. A similar suite of targets is observed if *sHAC1* is induced in mitotic cells (Pincus et al., 2014). Analysis of mRNA abundance for the 406 drug-induced UPR targets (Travers et al., 2000) indicated that most were unlikely to be Hac1 targets in pre-meiotic or mid-meiotic cells (Figure 3A). Only 32 transcripts were induced in pre-meiotic cells with timing that was consistent with control by Hac1 (turquoise box, Figure 3A), and only 58 were induced in mid-meiotic cells with timing that was consistent with the second wave of Hac1 synthesis (purple box, Figure 3A). Strikingly, these two sets of putative targets were entirely non-overlapping. This suggests that the well-defined stress-responsive UPR transcriptional program is tailored, and in two distinct ways, during the endogenous pre-meiotic and mid-meiotic inductions.

**Figure 3:**
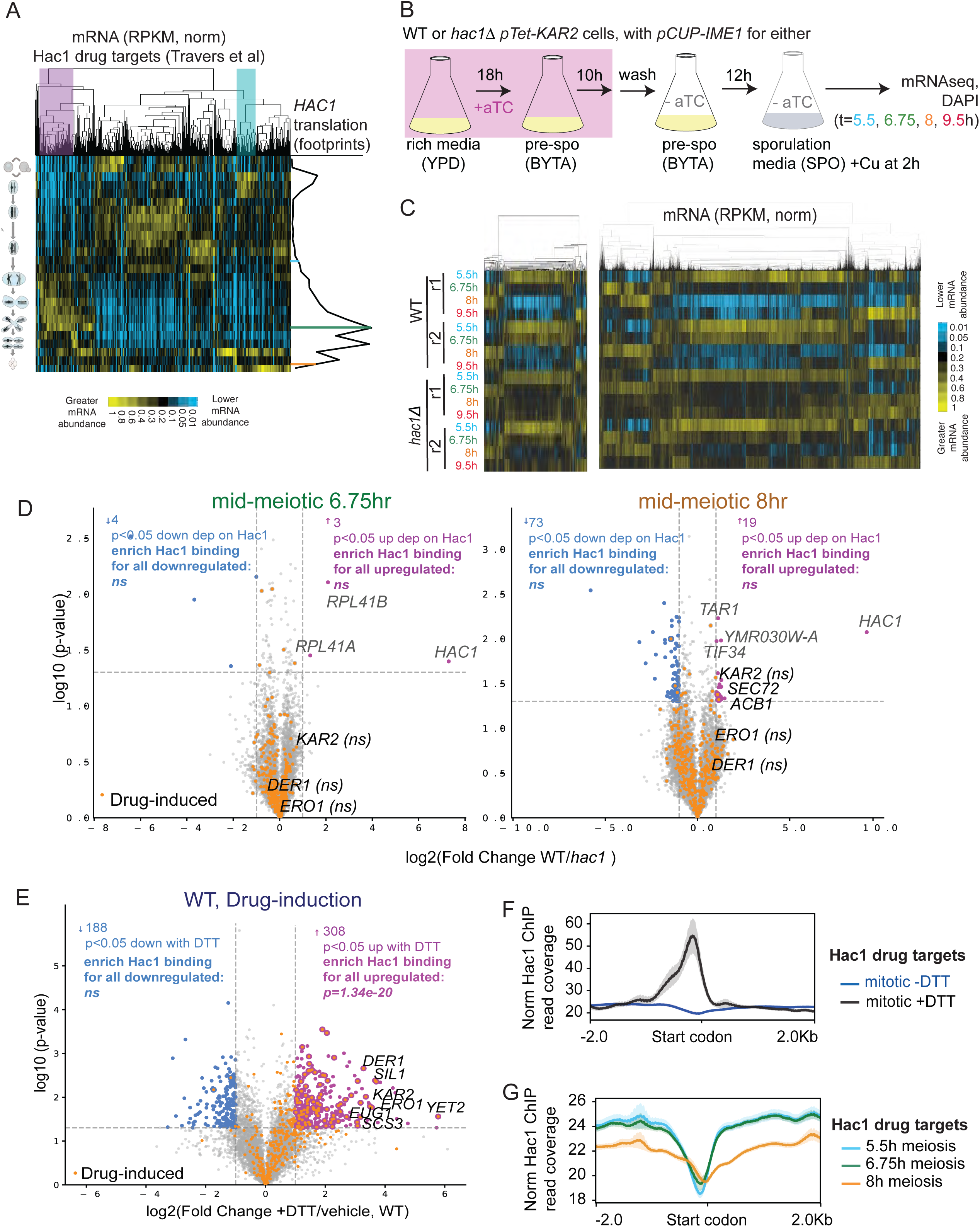
UPR activation in meiotic cells does not drive a canonical Hac1-driven transcriptional response. (A) mRNA-seq data from a previous study (Brar et al., 2012) for DTT- and Tunicamycin-defined UPR targets (Travers et al., 2000). Transcripts are subjected to hierarchical clustering, normalized over all timepoints. On the right, Hac1 translation from a parallel dataset is plotted. The purple box indicates the transcripts with abundance pattern that is consistent with mid-meiotic UPR induction, the turquoise box indicates transcripts with an abundance pattern that is consistent with pre-meiotic UPR induction. (B) Experimental scheme for globally assessing transcripts that depend on meiotic UPR induction. Cells were induced to enter meiosis hyper-synchronously through use of a *pCUP-IME1* allele and 50µM CuSO4 addition at 2h in SPO. (C) mRNA seq data from the experiment in (B) subjected to hierarchical clustering and normalized across all timepoints. On the left, only DTT- and Tunicamycin-defined UPR targets (Travers et al., 2000). On the right, all transcripts. (D) Volcano plots for 6.75hr in meiosis (left) and 8hr (right) display RNA-seq differential expression as log₂ fold change versus −log₁₀(p-value). Expression matrices were size-factor normalized and log₂(value + 1) transformed. Fold changes and p-values were calculated per gene using the mean of log₂-normalized replicates and Welch’s two-sample t-test, respectively. Genes with nominal p < 0.05 and |log₂FC| > 1 are colored magenta (upregulated dependent on Hac1) or blue (downregulated); all other genes are gray. Overlaid gene set is drug-induced targets (Travers et al. 2000) and are marked with orange center dots regardless of significance. No enrichment is seen among up or downregulated transcripts for genes with experimentally determined Hac1 promoter binding in matched conditions (File S2) as assessed by one-sided Fisher’s exact test. (E) Similar analysis as (D), but analyzing mitotic cells treated with DTT versus those that are mock treated. Highly significant enrichment among DTT-dependent upregulated genes for Hac1 binding in the promoter, as assessed by one-sided Fisher’s exact test. (F) Hac1 ChIP data comparing mitotic exponential cells treated with 4.5mM DTT or vehicle control. Metagene analysis for only DTT- and Tunicamycin-defined UPR targets (Travers et al., 2000), plotted is the average of two replicates. (F) Hac1 ChIP data comparing three meiotic timepoints. Metagene analysis for only DTT- and Tunicamycin-defined UPR targets (Travers et al., 2000), plotted is the average of two replicates. Hac1 signal is normalized by CPM.

We first assayed which transcripts are driven by the wave of Hac1 synthesis during mid-meiosis by growing *hac1Δ pTET-KAR2* cells in pre-meiotic media in the presence of aTC, then transferring them to sporulation media in the absence of aTC, using *pCUP-IME1* to achieve high synchrony (Berchowitz et al., 2013; Chia and van Werven, 2016). Under these conditions, meiosis occurs with similar synchrony and at a similar rate in *hac1Δ* and WT cells (Figure 3B, S1I, S1J). We collected replicate samples at several timepoints from WT cells and those lacking Hac1, including before (5.5h), during (6.75h, 8h), and after (9.5h) mid-meiotic UPR induction is normally seen. As a control, we also collected mitotic cells either grown in the absence or presence of 4.5mM DTT for 30 minutes. *HAC1* and *KAR2* mRNA behavior in WT controls confirmed the timing of endogenous UPR activation in this experiment (Figure S1K-N, 1A). Global mRNA comparisons from WT and Hac1-deficient meiotic timecourses revealed surprisingly similar patterns in both cases (Figure 3C, right; File S1). This was also true when the data were analyzed for just the 406 drug-induced UPR targets (Figure 3C, left). A few known Hac1 targets displayed extremely mild Hac1-dependent induction, specifically at 8h, including *ACB1, SEC72,* and *KAR2* (Figure 3D, S1O, S1P, S2A). None, however, displayed a difference in abundance that was consistent with meiotic induction by Hac1 in the same simple manner as is observed with drug-based UPR induction (ie. reduced in the absence of Hac1 at 6.75h and 8h; Figures 3D, S2A). In contrast, DTT treatment drove induction of the set of targets that was previously defined in drug-based and *sHAC1-*based UPR induction studies (enrichment of drug-defined targets p=1.4×10⁻³⁵, for sHAC1-induced targets p=3.7×10⁻⁷⁵; Figure 3E, S2A, Travers et al., 2000; Pincus et al., 2014). Thus, physiological mid-meiotic UPR output differs dramatically from stress-induced stimulation of the same pathway, with a minimal effect on gene expression.

### Robust Hac1 binding is only seen at the KAR2 promoter during mid-meiosis

Given the surprising lack of transcriptional changes caused by physiological UPR activation during meiosis, we performed chromatin IP (ChIP) experiments, using a tagged version of Hac1 that maintained its function in protecting cells from DTT treatment (Figure S1R). Samples analyzed included mitotic cells with and without DTT, as well as mid-meiotic cells at three timepoints prior to and during Hac1 induction (5.5h, 6.75h, 8h; as in Figure 3B, File S2). Analysis of the data yielded from mitotic cells with and without DTT treatment indicated that Hac1 ChIP was sensitive and specific. Metagene analyses revealed a strong and specific peak 5’ to start codons in DTT-treated cells that was more pronounced when just the 406 previously defined drug-responsive loci were analyzed (Figure 3F, S2B-C). Peak calling revealed Hac1 binding in the presence of DTT upstream of 286 CDSs, including many of the reported drug-induced UPR targets, and including well characterized targets *PDI1, KAR2, EUG1, ERO1,* and *HAC1* (File S2). Analysis of mitotic cells without DTT revealed no overall signal for Hac1 binding (Figure 3F), but peak calling revealed Hac1 binding at several loci, including *KAR2,* consistent with our previous finding that very low level UPR induction in mitotic conditions is important for maintaining expression of a subset of ER proteins, including Kar2 (Figure S2F, File S2; Bartolutti et al., 2025).

Parallel analysis of the mid-meiotic ChIP data revealed that a lack of Hac1 binding was seen, independently of whether the data for all genes were analyzed or if the subset of drug-induced targets were isolated for analyses (Figure 3G, S2D, S2E). At the 5.5h timepoint, only five Hac1 peaks were observed upstream of individual CDSs, and because we do not observe Hac1 splicing at this timepoint (Figure 1A, S1M, File S2) or mRNA induction of these targets, we consider these to represent background signal. At the 6.75h and 8h meiotic timepoints, only a single locus—*KAR2*—displayed a peak indicative of Hac1 binding in both replicates, and also mRNA induction consistent with Hac1 dependence, albeit mildly and only at 8h (Figure S2F). Four other ChIP peaks were observed (*ARG82/HMO1, YAP1802/FMP43, DCP2/MLS1, CPA1/RPA14*) but none of the proximal genes displayed regulation consistent with dependence on Hac1. These findings are consistent with a recent study revealing poor overlap between transcription factor binding and mRNA induction (Mahendrawada et al., 2025). Thus, together our data suggest that during endogenous mid-meiotic UPR induction, cells do not induce the type of broad transcriptional response that is characteristic of the drug-based UPR.

### Hac1 can drive drug-induced UPR targets in mid-meiosis when expressed highly

We were surprised at the low impact of Hac1 production during mid-meiosis on the transcriptional landscape (Figure 3C), given the robust Hac1-dependent transcriptional response seen with drug treatment of mitotic cells (Figure 3E; Travers et al., 2000), and reasoned that this could either be a result of the cellular context of mid-meiotic cells or the lower level of Hac1 expression seen under endogenous activation conditions. To distinguish between these possibilities, we examined the transcriptional impact of the UPR in mid-meiotic cells by collecting mRNA-seq samples at several timepoints with and without strong overexpression of *sHAC1,* comparing to mitotic samples with a similar strategy for Hac1 overexpression (Figure S3A-C, File S3-S4). For these experiments *sHAC1* mRNA was present in mitotic cells at a level >10-fold seen for *HAC1* (unspliced + spliced) in WT non-stress conditions, and in mid-meiotic cells at >2-fold the level of normal total *HAC1* mRNA (unspliced + spliced). Analyses of these data revealed robust and specific induction of most reported drug-induced or *sHAC1*-induced UPR targets in both mitotic and mid-meiotic cells (Figure S3B, S3C; enrichment of drug-defined targets p=7×10⁻⁷¹, for sHAC1-induced targets p=1×10⁻⁸⁵). This finding suggests that the lack of robust transcriptional changes associated with the mid-meiotic UPR, compared to the well-established drug-driven transcriptional program, is based at least in part on the lower levels of Hac1 with this physiological induction.

### UPR-deficient pre-meiotic cells display ER-associated reticulon foci and cannot enter meiosis

Both the transcriptomic data and analysis of meiotic progression for cells lacking UPR activity specifically within meiosis indicated that the mid-meiotic wave of endogenous UPR activation was not critical for maintaining the meiotic gene expression program (Figure S1E, 3C). However, UPR activity has also been reported to function in driving ER remodeling (Schuck et al., 2009), and we previously reported that dramatic changes in ER structure occur during meiosis, including the programmed collapse of most cortical ER (Otto et al., 2021). We therefore visualized ER morphology through a meiotic timecourse in cells with and without Hac1 by using a fluorescently tagged reticulon (Rtn1-GFP) as an ER marker. In the absence of Hac1, cells that were allowed to enter meiosis by *sHAC1* expression in pre-meiotic media displayed the characteristic changes in ER structure that typify normal meiotic progression (Figure 4A). During mid-meiosis, cortical ER largely collapses, and during gamete maturation, ER is again observed at the cell periphery. No differences between WT and UPR-deficient cells were evident (Figure 4A).

**Figure 4:**
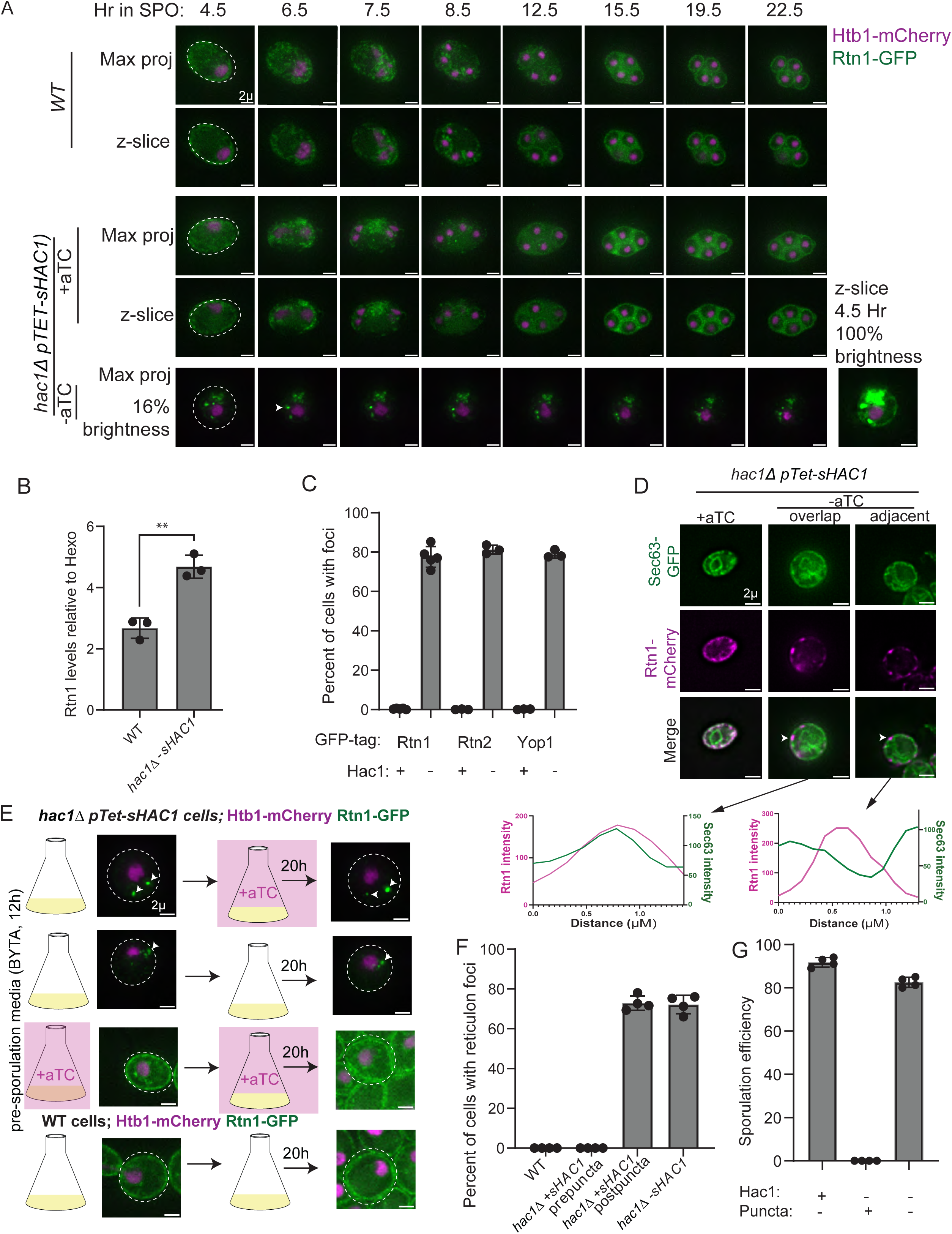
Pre-meiotic UPR-deficiency leads to irreversible reticulon focus formation. (A) Timelapse microscopy of WT cells, or *hac1Δ pTET-sHAC1* cells with or without aTC in SPO. Htb1-mCherry is shown in magenta, Rtn1-GFP in green. Note that brightness is not matched for foci containing cells for visualization purposes. One representative image at matched brightness to WT for reference is included. (B) Quantification of western blots probing for Rtn1-GFP normalized to Hexokinase quantification. Protein was isolated from cells grown in BYTA for 21 hours. n = 3 biological replicates. (C) *hac1Δ pTET-sHAC1* cells with or without aTC in pre-meiotic media, imaged for the presence or absence of reticulon foci. 300 cells were counted per strain per experiment, minimum of n=4 experiments. (D) *hac1Δ pTET-sHAC1* cells with or without aTC in pre-meiotic media. Rtn1-mCherry in magenta, Sec63-GFP in green. White arrowheads indicate two different locations featuring Rtn1-mCherry puncta that were analyzed with linescan analysis. Linescans of Rtn1 foci with arrows are below to demonstrate the overlapping versus adjacent colocalization with Sec63-GFP. (E) Imaging of *hac1Δ pTET-sHAC1* cells in pre-meiotic media. aTC is supplied at the indicated times. (F) Summary of imaging of WT or *hac1Δ pTET-sHAC1* cells, either without aTC addition or with aTC added prior to or after puncta form. 300 cells were counted per experiment, n=4 experiments. (G) Sporulation efficiency as scored by microscopy following 24 hours in sporulation media. 300 cells were counted per category per experiment, n=4 experiments.

In control cells that lacked Hac1 and that were not supplied with additional *sHAC1*, however, we observed bright Rtn1-GFP puncta within 80% of cells during pre-meiotic growth (Figure 4A, 4C). In these cells, normal cortical ER localization of Rtn1-GFP was still seen but the puncta were so bright that this signal could not be observed at exposure times that clearly show puncta (Figure 4A, compare 4.5h frame at right and left). We observed a higher overall level of Rtn1 protein in cells lacking Hac1 (Figure 4B, S4A), which may contribute to puncta formation. Puncta were never observed in WT cells or in UPR-deficient cells supplied with *sHAC1* in pre-meiotic media, which express Rtn1 at a level that is intermediate between WT and Hac1-deficient cells (Figure 4A-C). Similar foci were seen in cells expressing only *ire1-cLDΔ,* which were unable to splice *HAC1* in pre-meiotic or mid-meiotic stages but not in cells expressing the *bipless* and *R537* mutants, which display functional meiotic UPR at both stages (Figure 2D-2F, S4B, S4C). Bright puncta in pre-meiotic cells lacking Hac1 were also seen when the other yeast reticulon, Rtn2, or the reticulon-like Yop1 are visualized (Figure 4C, S4D, S4E). However, imaging of general ER marker Sec63, and all other ER proteins that we tested, did not reveal puncta under these conditions (Figure S4D, S4E).

Imaging cells carrying fluorescently tagged Sec63 and Rtn1 allowed us to determine that the reticulon foci either fully colocalized with ER regions containing Sec63 or were adjacent to it, consistent with their localization to the ER (Figure 4D, S4F). This result was supported by use of ERtracker (Figure S4G, S4H). Suggesting that these foci do not represent protein aggregates, we never observed puncta with PROTEOSTAT staining under these conditions (Figure S4I). Formation of these puncta was irreversible, as they could not be removed by expression of *sHAC1s* after their formation (Figure 4E, 4F). They were highly specific to the growth conditions in pre-meiotic media, as they were never observed in cells lacking *HAC1* that were grown in rich media (YPD), less nutrient rich synthetic media (SC) or the respiratory stimulating media YPG, which contains glycerol in place of glucose (Figure S4J, S4K). Pre-meiotic cells containing reticulon puncta never entered meiosis, whereas the minority of *hac1Δ* cells that did not display such puncta formed spores at a rate only slightly less than WT cells (Figure 4G).

### Pre-meiotic cells display non-canonical UPR-driven transcriptome remodeling

Based on the important role for Hac1 expression during pre-meiotic stages for both meiotic progression and normal ER morphology (Figure 1G, 4A), we sought to determine the transcriptional response of endogenous UPR activation prior to meiosis. We grew WT cells or cells lacking Hac1 (*hac1Δ pTET-KAR2*, in the absence of aTC) and collected replicate samples after 6h and 21h of pre-meiotic growth, representing log phase and saturated pre-meiotic growth, respectively (Figure 5A, S1Q). We also collected control samples in which pre-meiotic *hac1Δ pTET-KAR2* cells were grown in the presence of aTC, which rescues gamete formation (Figure 2B). Finally, we collected parallel Hac1 ChIP data for UPR-proficient cells (*3v5-HAC1*). In contrast to what we observed during mid-meiosis (Figure 3D), comparison of WT and UPR-deficient pre-meiotic cells revealed major overall differences in mRNA patterns for known drug-induced UPR target transcripts in pre-meiotic cells (Figure 5B). However, given that Hac1 splicing was present, and at a similar level in the early and late pre-meiotic growth (Figure S1Q), it was surprising that neither established drug-induced Hac1 targets nor *sHAC1*-induced targets (Travers et al., 2000; Pincus et al., 2014) showed higher abundance in WT cells at both pre-meiotic exponential and saturated growth stages, which would indicate transcription driven by Hac1 (Figure 5D, 5E, S2G). Instead, based on hierarchical clustering analysis, a cluster of 48 of these targets were higher in WT cells than in Hac1-deficient cells in early pre-meiotic growth, and a cluster of 29 in late pre-meiotic growth (Figure 5B). The early Hac1 pre-meiotic targets included well-characterized UPR targets and ER protein-encoding transcripts *KAR2* and *ERO1*, but their induction was very mild and not statistically significant based on analysis of replicate data (Figure 5D, S5A).

**Figure 5:**
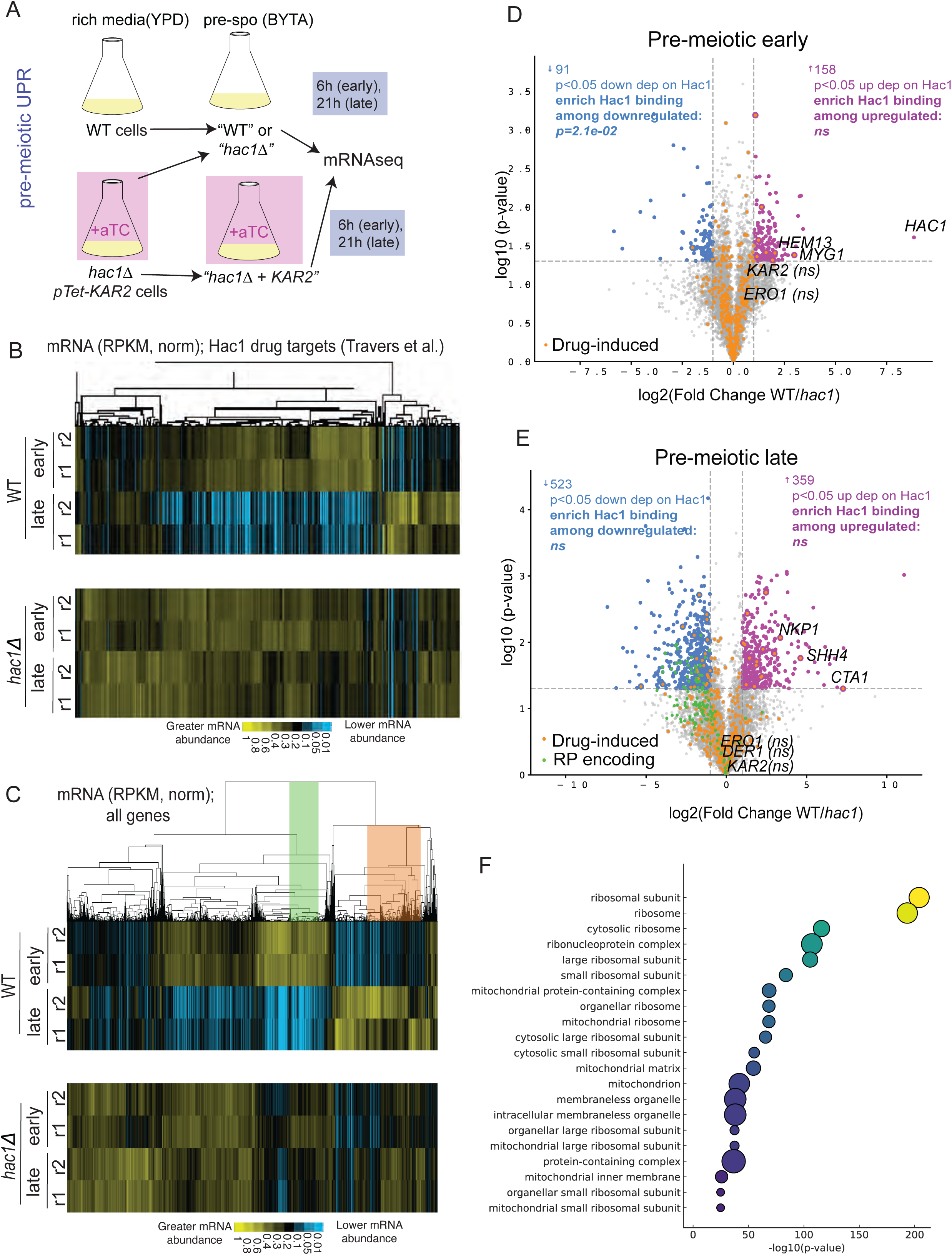
Pre-meiotic cells do not activate drug-induced UPR transcriptional targets. (A) Experimental scheme for globally assessing transcripts that depend on meiotic UPR induction. Cells were harvested at either 6h (early) or 21h in pre-meiotic media. (B) mRNA seq data from the experiment in (A) subjected to hierarchical clustering and normalized across all timepoints for only DTT- and Tunicamycin-defined UPR targets (Travers et al., 2000). (C) mRNA seq data from the experiment in (B) subjected to hierarchical clustering and normalized across all timepoints. The green and orange box indicate highly differentially expressed transcript patterns in the presence and absence of Hac1. (D) Similar analysis as in Figure 3D, except analyzing cells with and without Hac1 in a log-phase pre-meiotic (early) timepoint. Note that no enrichment is seen for experimentally determined Hac1 promoter binding in matched conditions among Hac1-dependent upregulated transcriptional targets but a mild enrichment is seen among downregulated transcriptional targets, potentially reflecting the type of regulation reported in (Mahendrawada et al., 2025) . (E) Similar analysis as in (D) but for the saturated (late) pre-meiotic timepoint. RP-encoding transcripts are colored as green dots, regardless of significance. (F) GO term analysis for cellular components for genes indicated by the green box in (C).

We observed major differences in the transcriptome of pre-meiotic cells, dependent on expression of Hac1 (Figure 5B-E, File S1). However, few established UPR targets are robustly induced in a Hac1-dependent manner in either pre-meiotic timepoint, and those that are induced are not those that are most highly induced by either DTT treatment or sHAC1 expression (eg. *CTA1, HEM13, MYG1, SHH4* rather than classical ER quality control factors; 5D, 5E, S2G, compare to 3E or S2A top). Although many sites of Hac1 binding were observed at these pre-meiotic timepoints, including upstream of *KAR2* and *ERO1* as assessed by ChIP peaks (Figure S2L), there was no enrichment of these binding sites among upregulated targets in either pre-meiotic timepoint (Figure 5D, 5E; S2H-K; File S2). Hierarchical clustering of the full dataset revealed two major transcript groups of interest. The first contained 418 transcripts that were much higher in early pre-meiotic cells and much lower in late pre-meiotic cells when Hac1 was present (green box, Figure 5C). The second cluster contained 734 transcripts that were higher in late pre-meiotic cells with Hac1 (orange box, Figure 5C). The late-induced transcripts were enriched for functions in catabolism and peroxisome function (Figure S5C). The early-induced/late-repressed group were strongly enriched for roles in translation and included ribosomal protein (RP) encoding genes and those involved in ribosome biogenesis (Figure S5B, 5F).

### Pre-meiotic UPR induction leads to repression of translation

Analysis of the saturated (late) pre-meiotic timepoint revealed nearly universal downregulation of RP-encoding transcripts in a Hac1-dependent manner (Figure 5E, 6A). Expression of *sHAC1* in mitotic cells resulted in reduced transcript abundance for RP genes (Figure 6B), as did treatment with common UPR trigger DTT, in a manner dependent on Hac1 (Figure 6C). Proteomic analysis of pre-meiotic cells with and without Hac1 revealed reduced protein levels for RPs, mirroring the effect observed for their transcripts (Figure 6D; File S5). Together these data indicate the UPR has a general role in reducing RP gene expression in yeast. To test whether these reductions in expression of proteins that support translation led to a reduction in bulk translation, we performed puromycin-incorporation experiments in pre-meiotic cells, at exponential (early) and saturated (late) stages in cells expressing or not expressing Hac1. Consistent with the trends in RP gene expression, we observed a reduction in bulk translation in WT cells at late pre-meiotic stages, as cells prepare to enter meiosis (Figure 6E, S5F). Moreover, pre-meiotic cells lacking Hac1 are defective at this repression. Reduced RP gene expression has been observed as a secondary effect of defective growth (Brauer et al., 2008), and expression of *sHAC1* has been reported to slow cell growth (Pincus et al., 2014). However, this cannot explain the effect that we observe in pre-meiotic cells, as cells deleted for *HAC1* grow more slowly under these conditions than their WT counterparts (Bartolutti et al., 2025).

**Figure 6:**
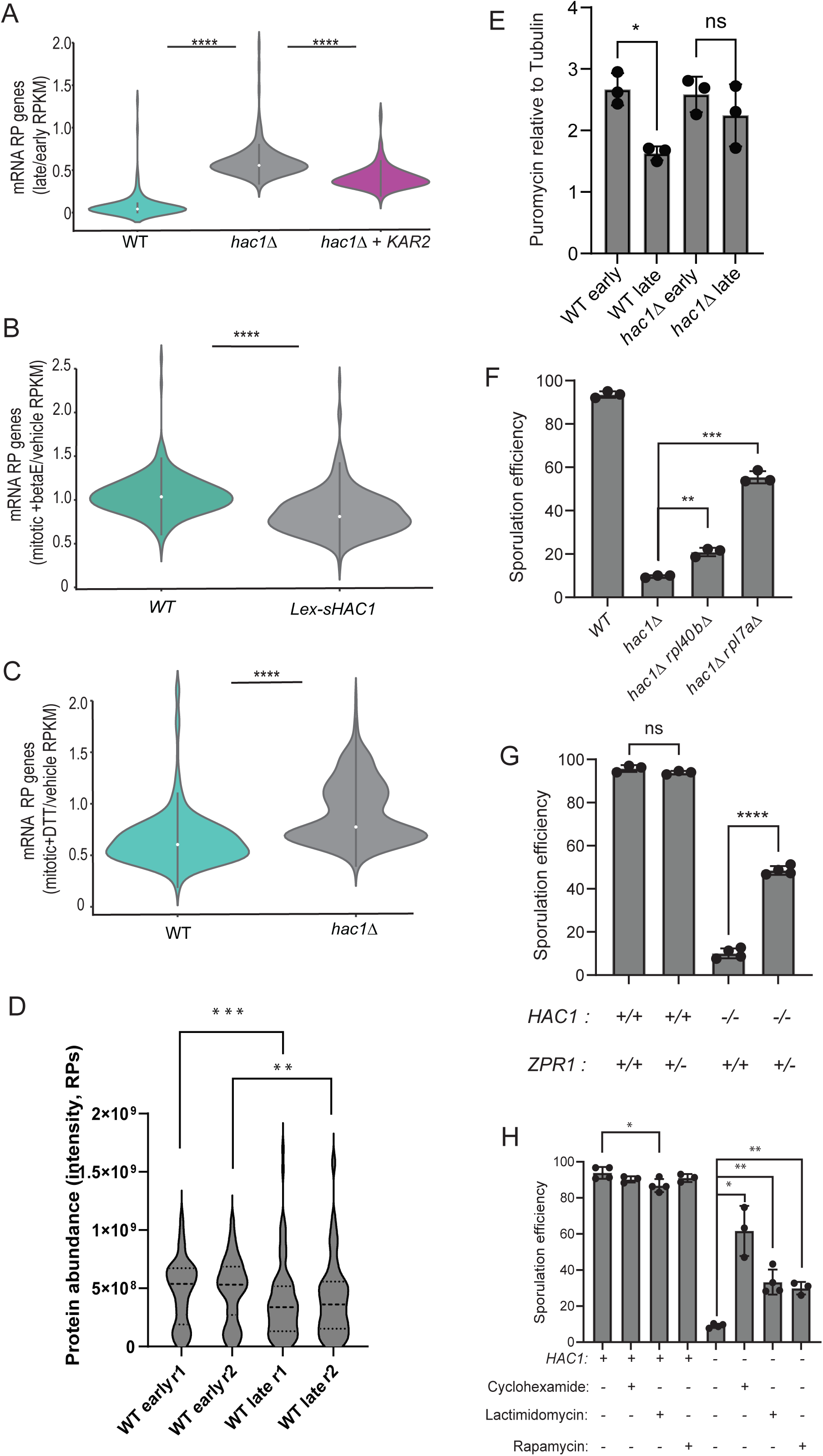
Pre-meiotic UPR activity supports meiotic entry by repressing translation (A) Plot of the ratio of mRNA RPKM (reads per kilobase million) at late/early pre-meiotic timepoints for all ribosomal protein (RP) genes for the conditions schematized in Figure 5A and shown in Figure S6D. (B) Plot of the ratio of mRNA RPKM for vegetative cells with and without beta-estradiol-driven *sHAC1* expression for all RP genes in WT. (C) Plot of the ratio of mRNA RPKM for vegetative cells with and without 4.5mM DTT treatment for all RP genes in WT or *hac1Δ* cells. (D) Plot of the ratio of protein quantification by label free mass spectrometry intensity values for late/early pre-meiotic timepoints in WT cells. Included are protein values for all quantified RPs for two biological replicates. (E) Translation activity as assessed by western blotting for puromycin incorporation of WT or *hac1Δ* cells grown in pre-meiotic media (BYTA) at log (early) or saturated (late) stages. Data represent three biological replicates, blot shown in Figure S5F. Puromycin signal was normalized to tubulin signal. (F) Sporulation efficiency as scored by microscopy following 24 hours in sporulation media. 300 cells counted per strain/condition per experiment, n=3. (G) Sporulation efficiency as scored by microscopy following 24 hours in sporulation media. 300 cells counted per strain/condition per experiment, minimum of n=3. (H) Sporulation efficiency as scored by microscopy following 24 hours in sporulation media. 300 cells counted per strain/condition per experiment, minimum of n=3. Significance was determined from unpaired t-test with Welch’s correction. P-value significance is **** < 0.0001, *** < 0.001, ** < 0.01., * < .05.

### Reducing translation rescues gamete production in UPR-deficient cells

Reduced bulk translation is seen as an important adaptive feature of many stress responses, including the integrated stress response, but has not been reported as a consequence of the Ire1 branch of the UPR. We wondered whether repression of translation via downregulation of RP genes alone could drive meiotic progression in pre-meiotic cells lacking Hac1. We leveraged two previously validated mutant strain backgrounds (*rpl40bΔ* and *rpl7aΔ)* to reduce ribosome number and bulk translation to 64% and 51% of WT levels, respectively (Cheng et al., 2019). These mutant cells grow more slowly mitotically than WT controls, as expected, with a 23% and 50% increase in doubling time, respectively (Cheng et al., 2019). Remarkably, however, when combined with deletion of *HAC1,* these mutants rescued the defect in gamete production seen in cells deleted for *HAC1* alone, proportional to their ability to reduce translation (Figure 6F). We also analyzed *hac1Δ* cells that were heterozygous for *ZPR1*, a gene encoding an essential chaperone that facilitates folding and function for the translation elongation factor eEF1A (Sabbarini et al., 2023), again noting a rescue in spore formation (Figure 6G). This rescue appears to be independent of activation of Hsf1, which is observed in cells lacking Zpr1 function, as classic Hsf1 targets are not upregulated (S5G).

Silencing of the pre-meiotic UPR in WT cells corresponds to the timing at which bulk translation increases during meiotic entry, despite a transfer to media that is more nutrient poor than pre-meiotic media (Brar et al., 2012). Together with the data above, this suggests that pre-emptive silencing of translation may be preparatory to enable meiosis. To test whether this might be the case, we used several pharmacological strategies for inhibiting translation, including treatment with low concentrations of either rapamycin, which indirectly reduces translation through Tor repression; or cycloheximide or lactimidomycin, which inhibit elongating and post-initiation ribosomes, respectively. Remarkably, in all cases we observed an increase in the ability of cells lacking Hac1, but not WT counterparts, to enter and complete meiosis (Figure 6H). These data suggest that the repression of translation that occurs as a consequence of UPR activity is a physiological pre-meiotic role for the UPR, serving a preparatory role in supporting meiotic entry.

### Chaperone overexpression or translation repression prevent the formation of reticulon foci in UPR-deficient cells

Having identified either chaperone expression or reduced translation as capable of suppressing the defect in meiotic success seen in UPR-deficient cells, we next tested whether either of these conditions impacted the presence of reticulon puncta that we observed in the absence of Hac1 in pre-meiotic cells (Figure 4E). Indeed, expression of Kar2 in *hac1Δ* cells prevented formation of these ER puncta, as did expression of Rot1, which was also able to rescue meiotic progression in *hac1Δ* cells (Figure 2B, 7A, 7B). Remarkably, reduction of bulk translation, through treatment of cells with either cycloheximide, lactimidomycin, or rapamycin — all conditions that rescue sporulation in UPR-deficient pre-meiotic cells — also prevented puncta formation (Figure 7C, 7D). These data indicate that either insufficient ER folding capacity or too much overall protein production leads to ER dysfunction and prevent meiosis in the absence of the UPR. Indicating that reticulon puncta may be indicative of defects in ER quality, expression of the cortical ERphagy adaptor Atg40 in pre-meiotic conditions (in which general autophagy factors are present) prevented the presence of reticulon puncta in pre-meiotic cells (Figure 7E, 7F). We observe an increase in autophagy of Rtn1 in Hac1-deficient cells, as assessed by the amount of free GFP in cells carrying Rtn1-GFP (Figure 7G, S5H), further supporting reduced ER quality in pre-meiotic cells lacking UPR support. Together, these data support the model that defects in ER folding are associated with aberrant ER protein structures and an inability of cells to enter meiosis.

**Figure 7:**
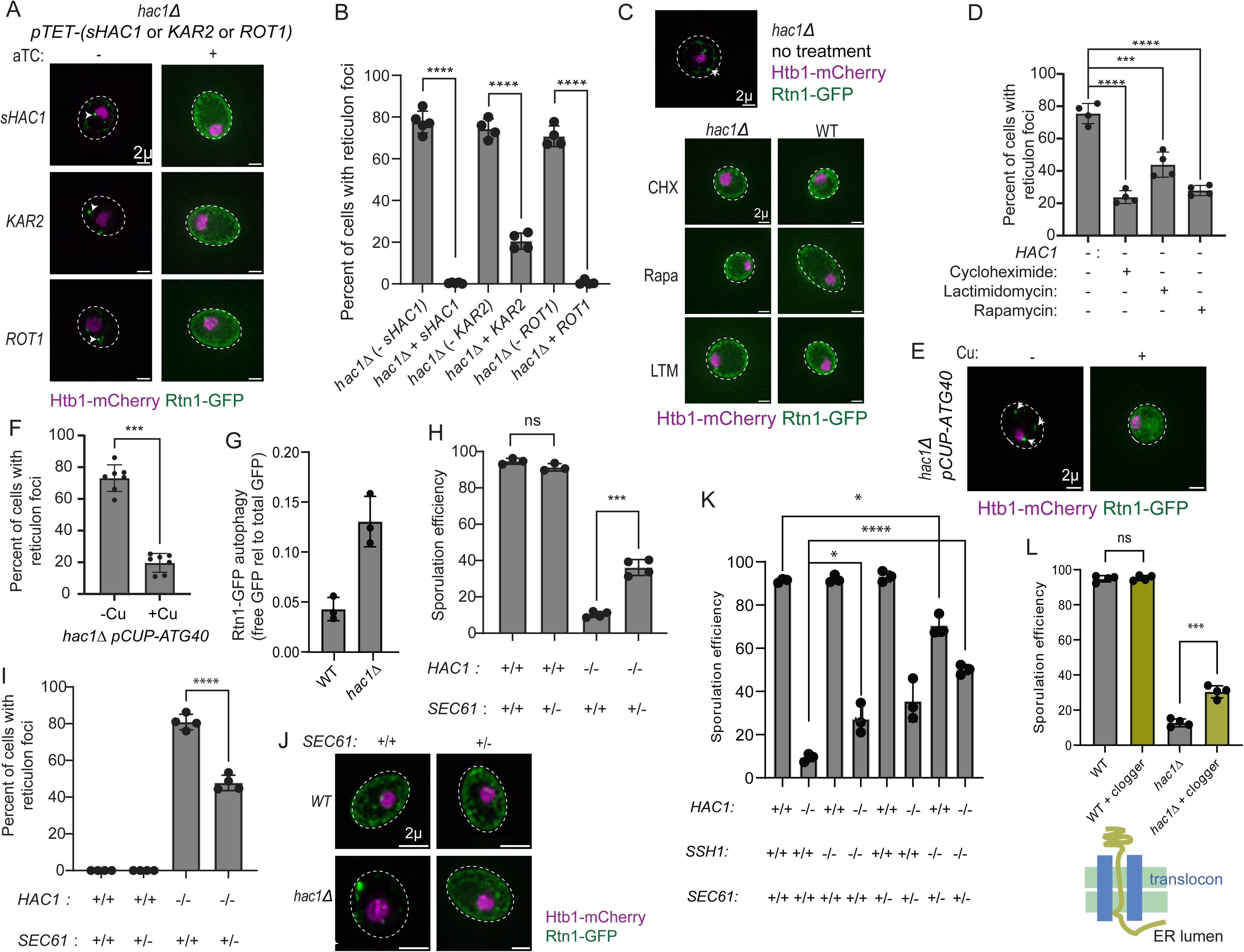
Pre-meiotic reticulon foci in UPR-deficient cells are prevented by chaperone expression, translation inhibition, and reduced ER translocation capacity. (A) Imaging of Rtn1-GFP morphology in *hac1Δ* cells supplemented with either *sHAC1*, *KAR2,* or *ROT1.* (B) Quantification of imaging as in (A). 300 cells counted per strain/condition per experiment, minimum of n=4. (C) Imaging of Rtn1-GFP morphology in *hac1Δ* cells treated with 50ng/mL cycloheximide (CHX), 100nM rapamycin (Rapa), or 30nM lactimidomycin (LTM) for 18 hours. (D) Quantification of imaging as in (C). 300 cells counted per strain/condition per experiment, n=4. (E) Imaging of Rtn1-GFP morphology in *hac1Δ* cells carrying *pCUP-ATG40* and either treated or not treated with 50µM CuSO4 for 18 hours. (F) Quantification of imaging as in (E). 300 cells counted per strain/condition per experiment, n=4. (G) Quantification of autophagy of Rtn1-GFP in WT and *hac1Δ* saturated pre-meiotic cells as assessed by the fraction of GFP signal in free versus total (full-length plus free) bands. Three biological replicates were analyzed, 1 representative blot in Figure S5H. (H) Sporulation efficiency as scored by microscopy following 24 hours in sporulation media. 300 cells counted per strain/condition per experiment, minimum of n=3. (I) Quantification of imaging as in (H). 300 cells counted per strain/condition per experiment, n=4. (J) Imaging of Rtn1-GFP morphology in *hac1Δ pTET-sHAC1* cells treated with or without 250ng/mL aTC and carrying either WT *SEC61* loci or heterozygously deleted for one copy of *SEC61.* (K) Quantification as in (H) for genotypes shown. n=3. (L) Either WT or *hac1Δ* cells were induced to express a translocon “clogger” construct, as described in (Ast et al., 2016) and schematized below. Sporulation efficiency was assessed as in (H). n=4. Significance was determined from unpaired t-test with Welch’s correction. P-value significance is **** < 0.0001, *** < 0.001, ** < 0.01., * < .05.

### Reduction in ER translocation bypasses the need for UPR activity to support meiosis

Why would a reduction in translation result from UPR activation and how could this be beneficial to ER integrity? We previously showed that meiotic cells undergo a profound remodeling of the ER during meiosis, including a near-complete collapse of the cortical ER from the plasma membrane, which could impact ER folding capacity (Otto et al., 2021). Moreover, we recently found that ER protein folding is of great importance to cellular fitness even in mitotic cells in the absence of stress (Bartolutti et al., 2025). This conclusion was based on data including that UPR-deficient mitotic cells often lost chromosome V (if diploid) to maintain fitness. The fitness advantage offered to UPR-deficient diploid mitotic cells by loss of chromosome V could be recapitulated by heterozygous expression of Sbh1, Sbh2, and Irc22, three factors related to ER translocation that are encoded on this chromosome. Intriguingly, these *hac1Δ* aneuploid cells displayed reduced reticulon puncta relative to *hac1Δ* controls and were able to complete meiosis at a markedly increased level (Figure S5J, S5K). *hac1Δ* cells that were also heterozygous for *sbh1Δ, sbh2Δ,* and *irc22Δ* similarly displayed increased capacity to undergo meiosis (Figure S5L), as did cells heterozygous for *sec61Δ*, a gene encoding an essential translocon component (Figure 7H). Importantly, these cells all also displayed a corresponding decrease in reticulon foci (Figure 7I, 7J). This rescue of sporulation was exacerbated by simultaneous reduction in Sec61 and deletion of the gene encoding its homolog Ssh1 (Figure 7K). To test whether these effects are mediated by reduction in ER translocation, we used a complementary approach based on a translocon “clogger” protein, with the natural ER protein Pdi1 fused to a rapidly folding stable C-terminal domain trapping the protein mid-translocation (Ast et al., 2016). Remarkably, expression of this clogger protein partly rescues sporulation in cells lacking Hac1 (Figure 7L). Together, these results suggest that decreased ER folding burden achieved by either reduced translocation, increased chaperone expression, or reduced translation can compensate for lack of a functional UPR in promoting gametogenesis.

Our data support the model that pre-meiotic cells are sensitive to deficiencies in ER folding capacity and that ER folding capacity during such stages is normally supported by endogenous activation of UPR activity. However, our data suggest that the output of the UPR under these conditions is markedly different than the canonical output, which includes a large suite of transcriptional targets related to ER quality control. In pre-meiotic conditions, the principle transcriptional changes instead appear to be related to reducing the load of new proteins produced. Together, our data suggest that in the absence of UPR activity in pre-meiotic cells, ER folding capacity is overwhelmed due to an increased load of proteins translocated into the ER. This state leads to aberrant ER morphology and an inability of cells to enter meiosis.

## Discussion

The Ire1 branch of the UPR is highly conserved and is important in embryogenesis, differentiation, and disease states (Reimold et al., 2001, 2002; Madden et al., 2019). These known critical roles of the UPR are endogenously driven rather than activated in response to exogenous triggers. Much is known about drug-induced UPR signaling but much less is known about physiological activation. Here, we report two UPR activation events associated with meiosis in budding yeast. The first occurs prior to meiotic entry and is required for it. This physiological UPR activation event diverges in several ways from what has been seen as part of the well-studied reactionary drug response. It does not induce the canonical Hac1 targets that are induced in response to DTT or tunicamycin treatment, and instead broadly represses translation, which has not been previously attributed to the Ire1 branch of the UPR. This pre-meiotic UPR activation is required to prevent formation of aberrant ER protein structures which are associated with the inability of cells to enter meiosis. Either expression of the ER chaperone Kar2, or reduction in translation can both rescue meiotic entry in UPR-deficient cells, although Kar2 expression is not naturally robustly induced by the pre-meiotic UPR, whereas reduced translation is. Both also prevent formation of the reticulon foci. These features of the physiological meiosis-associated UPR diverge dramatically from the response defined by drug-induction, highlighting the importance of studying physiological outputs of the UPR, a lesson that is also applicable to other stress-responsive pathways.

One of the most fascinating results of UPR deficiency in pre-meiotic cells is the formation of concentrated regions of reticulon localization. These foci of reticulons include all three yeast reticulon and reticulon-like proteins and colocalize with general ER markers but do not include concentration of any other ER proteins that we have tested thus far. Once formed, they appear to be irreversible, and when they are present, cells are not capable of entering meiosis. Reduced translation or reduced expression of Sec61 prevent focus formation in a UPR-deficient background, as does expression of Kar2, suggesting that reticulon focus formation is related to the dysfunctional ER folding environment in pre-meiotic cells lacking the UPR. These reticulon foci form specifically in pre-meiotic cells, but not in other UPR-deficient conditions that we have examined, and they can be prevented by overexpression of the ERphagy receptor Atg40. Suggesting that they represent defective or stressed ER, similar foci have recently been reported to form in response to harsh drug-based ER stressors in mitotic cells (Platzek et al., 2025). Further exploration of their nature, including whether they are sites of autophagy, and how they prevent meiosis or whether they are themselves a byproduct of a toxic process is warranted.

The second UPR activation event associated with meiosis occurs concomitant with the meiotic nuclear divisions in mid-meiosis. Both transcriptomic and ChIP analyses indicate that despite the presence of *HAC1* mRNA splicing and Hac1 protein production, this UPR does not drive a broad transcriptional program, instead very mildly and transiently boosting a few targets, including *KAR2,* in mid-meiosis. Although Kar2 overexpression can compensate for loss of the UPR in pre-meiotic cells, it is not robustly induced by Hac1 in pre-meiotic cells, and mid-meiotic UPR induction is dispensable for meiotic success, making the presence of this robust, physiological UPR induction perplexing. Overexpression of Hac1 in meiosis is capable of activating most of the targets that are activated in response to drug treatment, suggesting that lower level of Hac1 in meiosis compared to in response to drugs is part of the explanation for the lack of a broad transcriptional program driven by the mid-meiotic UPR. Also given the few targets directly activated in pre-meiotic cells, it seems likely that the type of major transcriptional response previously thought to be a core result of Ire1 activation may not be the functional output of physiological UPR activation. This is reminiscent of a recent study of the heat shock factor Hsf1, which reported its role in activating a narrower transcriptional program than previously thought (Solís et al., 2016).

Our data indicate that this role for the UPR in repressing translation-associated factors (including ribosomal proteins and ribosome biogenesis factors) is not unique to pre-meiotic activation and also occurs to a lesser degree as part of the drug-induced Hac1-dependent UPR, as well as in response to expression of *sHAC1* under unstressed conditions. This highlights a previously unclear parallel between the UPR in yeast and metazoans, as signaling through PERK and the RIDD function of Ire1 both help to reduce translation (Harding et al., 2000; Hollien and Weissman., 2006; Walter and Ron, 2011). An explanation for RIDD has been that it is a way for cells to reduce the load of proteins being translocated into the ER under conditions in which ER folding capacity is overtaxed. The ability of reduced expression of translocation factors, most notably Sec61, to rescue the need for pre-meiotic UPR activity is consistent with the model that overtaxed ER folding capacity is a problem for pre-meiotic cells in the absence of the UPR. Strikingly, even clogging the translocon can rescue sporulation in UPR-deficient cells. We do not know how the pre-meiotic UPR drives down translation. It may be indirect, but we also note that Hac1 binding can be detected upstream of the CDS for *DOT6,* a gene encoding a repressor of ribosome biogenesis factors that is highly regulated in response to cellular nutritional status (Kusama et al., 2022; Figure S5M). We also observe upregulation of *DOT6* mRNA in pre-meiotic cells that is dependent on the presence of Hac1 (File S1). It appears that Kar2 expression may mildly rescue RP expression in pre-meiotic UPR-deficient cells, but this is not likely the natural mechanism by which translation is modulated, given the lack of robust *KAR2* induction normally seen at this time. The specific mechanism by which the Ire1-Hac1 UPR downregulates translation is a fascinating area of future research.

Why reduced translation (particularly of translocated ER proteins) is beneficial to cells preparing to enter meiosis is an interesting area for future study. One hypothesis relates to the dramatic ER remodeling that occurs during meiosis, which includes restructuring of the cortical ER early in meiosis into cable-like structures and culminates with detachment of most cortical ER from the plasma membrane (Otto et al., 2021). It may be that these restructuring events limit ER folding capacity and that entry into meiosis depends on reduced overall protein production to allow these critical ER remodeling events to occur. Alternatively, it may be that a reduction in translation simply prepares cells to enter meiosis with a clean slate, enabling synthesis of the many specific waves of proteins required for this complex developmental process in a synchronous way (Brar et al., 2012). Whatever the reason, the events that we observe to be supported by the pre-meiotic UPR - reduced translation and maintenance of normal reticulon localization – are associated with successful meiotic entry. This type of preparative role for the UPR, as part of a natural induction, is distinct both conceptually and mechanistically from the type of reactive transcriptional response that occurs in response to drug treatment. It will be interesting to see if these distinctions are shared by internally driven metazoan UPR activation events, including during embryogenesis and B-cell differentiation (Reimold et al., 2000, 2001).

## Acknowledgements

We thank Nick Ingolia, Michael Rapé, and members of the Brar and Ünal labs for feedback on this manuscript. We thank Maya Schuldiner for generously sharing the translocon clogger construct. This work was supported by NIH funding to GAB (R35GM134886), NIH funding to MJ (R35GM152258), as well as NIH (R01AG071801) and Astera Institute to EÜ. CB was funded by T32GM008295 and T32GM148378. AB was funded by an NSF predoctoral fellowship (DGE2146752) and T32GM132022.

---

**Figure S1:**
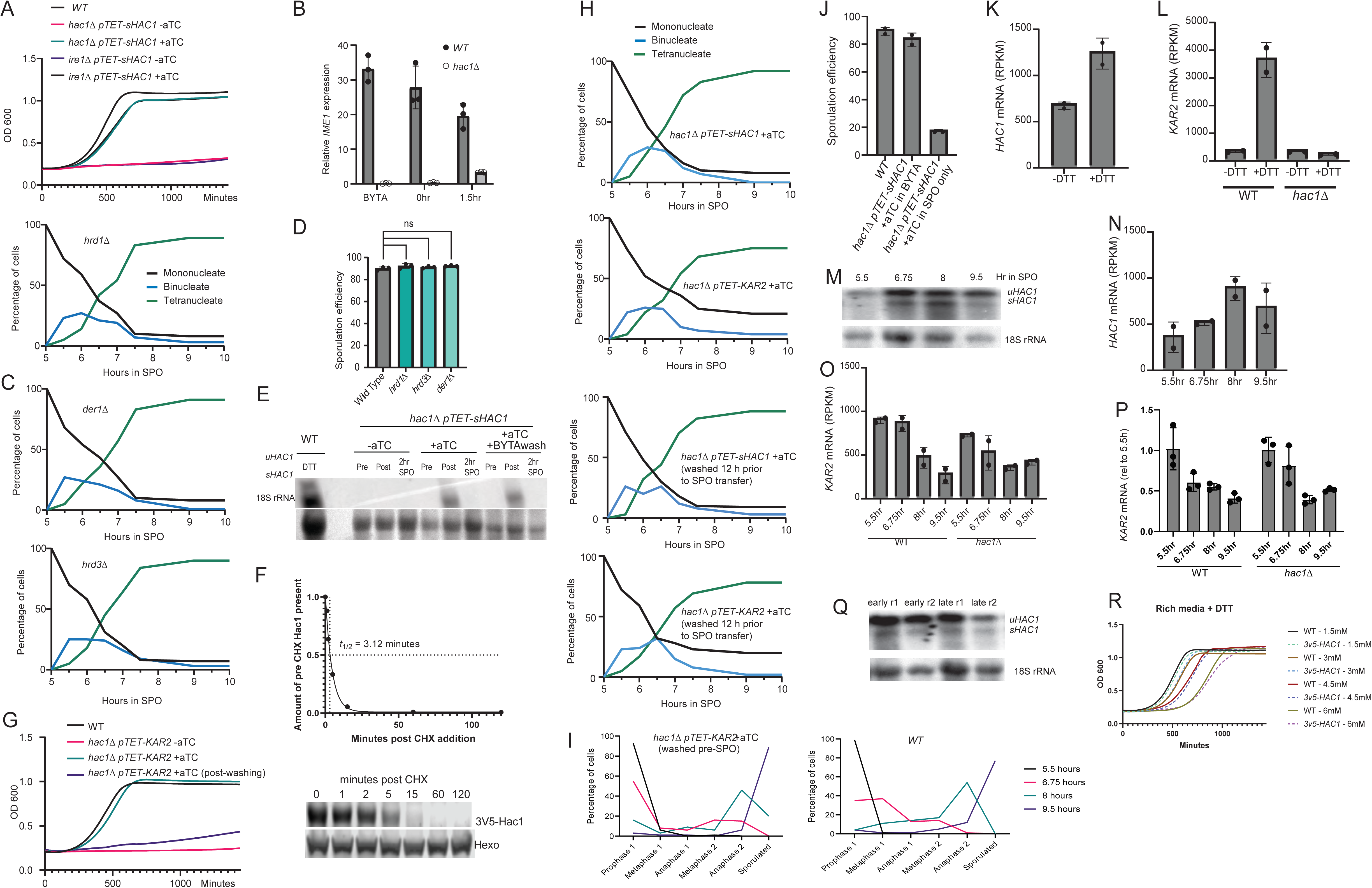
Pre-meiotic UPR activity supports normal meiotic progression (A) WT, and *hac1Δ pTET-sHAC1* or *ire1Δ pTET-sHAC1* cells either treated with 250ng/mL aTC or not treated with aTC growth in the presence of 1.5mM DTT in YPD. Cells were grown in YPD to saturation overnight before back dilution (B) *IME1* expression, relative to *ACT1*, as determined by RT-qPCR in either pre-meiotic media (BYTA), immediately after transfer to sporulation media, or after 1.5h in sporulation media, biological replicates n=3. . (C) Meiotic progression scored by Mono-, bi-, and tetra-nucleate cells assessed by DAPI staining. 100 cells counted per timepoint and strain. (D) Sporulation efficiency as scored by microscopy following 24 hours in sporulation media. 300 cells counted per strain/condition per experiment, minimum of n=3. (E) Northern blotting to detect unspliced (*u)* and spliced (*s) HAC1* mRNA in *hac1Δ pTET-sHAC1* cells without aTC, with aTC, or with aTC which was washed after 10 hours of growth. All strains were washed after 22 hours of total growth in BYTA before media was switched to SPO. (F) Hac1-3V5 protein abundance as determined by western blotting on the right and quantified on the left. Hac1-3V5 was overexpressed via addition of 5nM beta estradiol at the start of growth in BYTA media to drive the lex-sHac1-3v5 construct. 100ug/mL cycloheximide was added after 21 hours of growth in BYTA and protein samples were collected. Immunoblot (below) is for V5 and Tubulin with quantification done by normalizing the V5 intensity for each lane by the corresponding Tubulin intensity. Data was fitted to a one phase exponential decay model using GraphPad Prism software and t_1/2_ was calculated from this model. (G) WT or *hac1Δ pTET-KAR2* cells either treated with 250ng/mL aTC, not treated with aTC, or treated with aTC for 10 hours, after which it was washed out for 12 hours. All conditions were washed at end of BYTA growth before growth in YPD +1.5mM DTT was started. Growth is assessed by OD600 measurement of culture density.(H) Meiotic progression scored by Mono-, bi-, and tetra-nucleate cells assessed by DAPI staining. 100 cells counted per timepoint and strain. (I) Meiotic progression for experiment in Figure 3B,as scored by spindle morphology with tubulin IF. 200 cells counted per timepoint per strain background. (J) Sporulation efficiency as scored by microscopy following 24 hours in sporulation media. 250ng/mL of aTC was added to *hac1Δ* cells carrying *pTET-sHAC1* either at the same time as media was switched to BYTA or when media was switched to SPO. 300 cells counted per strain/condition per experiment, minimum of n=4. (K) *HAC1* RPKM values plotted from mRNAseq data for mitotic WT cells -DTT and +DTT treatment. (L) *KAR2* RPKM values plotted from mRNAseq data for mitotic –DTT and +DTT samples. (M) Northern blot probed for *HAC1* as in Figure 1A. RNA samples from samples used for meiotic mRNAseq in Figure 3. One representative replicate shown. (N) *HAC1* RPKM values plotted from mRNAseq data for meiotic samples. (O) *KAR2* RPKM values plotted from mRNAseq data for meiotic samples with or without Hac1. (P) *KAR2* relative mRNA abundance measured by qPCR analysis. RT was done with the same RNA used for mRNAseq. qPCR was normalized to actin as a loading control then all samples were plotted as fold change compared to their respective 5.5hr average. n=3 qPCR technical replicates plotted. (Q) Northern blot probed for *HAC1* as in Figure 1A. RNA samples from samples used for BYTA mRNAseq in Figure 5. (R) WT and *3v5-HAC1* cells grown in the presence of various concentrations of DTT in YPD. Cells were growth in YPD to saturation overnight before back dilution. Significance was determined from unpaired t-test with Welch’s correction. P-value significance is **** < 0.0001, *** < 0.001, ** < 0.01., * < .05.

**Figure S2:**
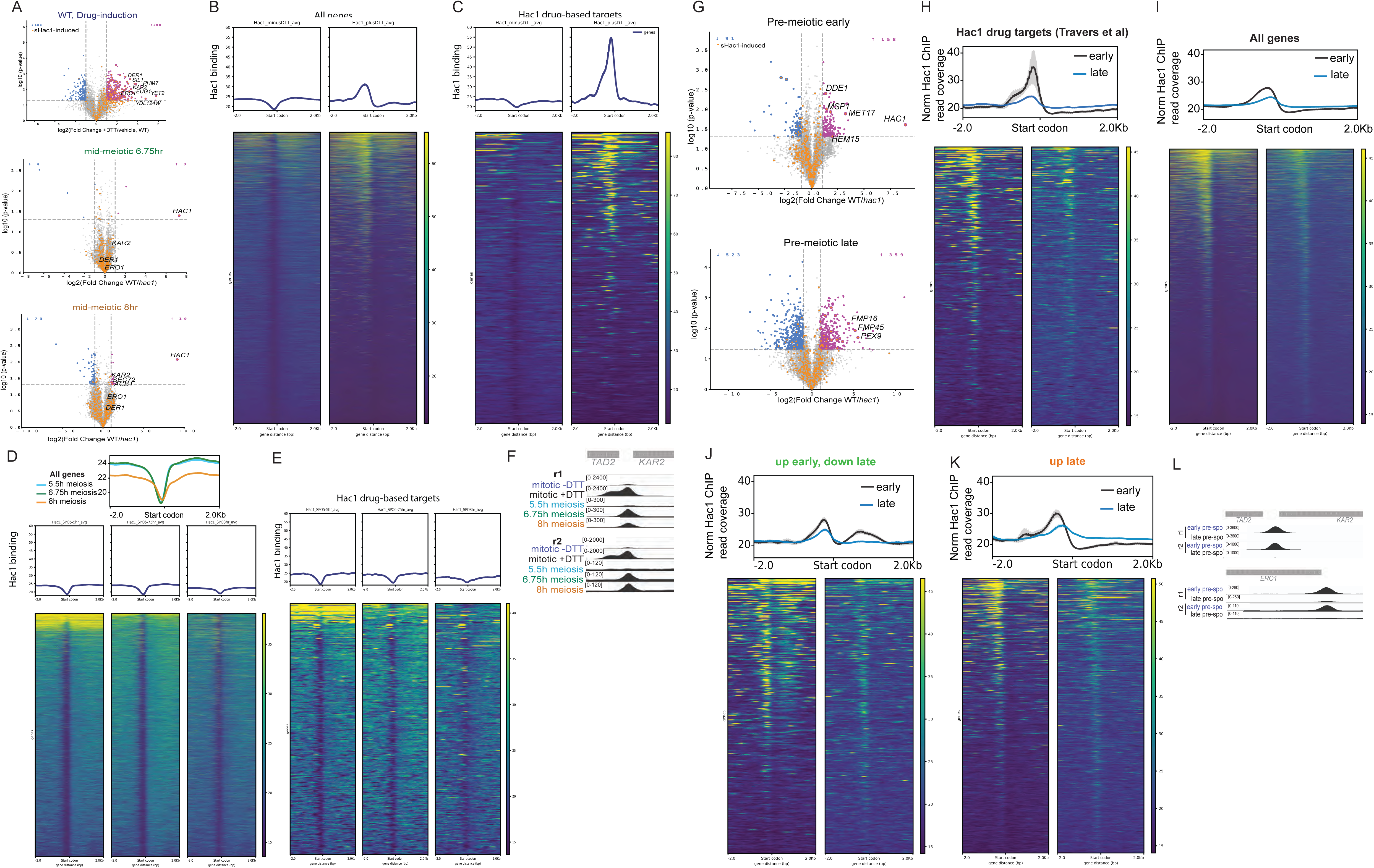
Hac1 binding cannot explain the patterns of mRNA expression for most meiotic and pre-meiotic UPR targets (A) Similar analysis as in Figure 3D and 3E, but overlaying genes with orange dots that are induced by *sHAC1* in (Pincus et al. 2014). (B) At top are metagene analysis of Hac1 binding around all CDS start codons in mitotic cells either without or with DTT, at bottom are traces of each gene, with colorimetric scale indicating amount of Hac1 binding, with yellow as high and blue as low. (C) At top are metagene analysis of Hac1 binding around CDS start codons for drug-induced UPR targets in mitotic cells either without or with DTT, at bottom are traces of each gene, with colorimetric scale indicating amount of Hac1 binding, with yellow as high and blue as low. (D) At top are metagene analysis of Hac1 binding around all CDS start codons in meiotic cells after 5h, 6.75h, or 8h in SPO. Metagene analysis for all annotated CDSes, plotted is the average of two replicates. Hac1 signal is normalized by CPM. At bottom are traces of each gene, with colorimetric scale indicating amount of Hac1 binding, with yellow as high and blue as low. (E) At top are metagene analysis of Hac1 binding around all CDS start codons for drug-induced UPR targets in meiotic cells after 5h, 6.75h, or 8h in SPO. (F) Hac1 ChIP signal around the *KAR2* locus with two replicates (r1 and r2) plotted. (G) Similar analysis as in Figure 5D and 5E, but overlaying genes with orange dots that are induced by *sHAC1* in (Pincus et al. 2014). (H) At top are metagene analysis of Hac1 binding around all CDS start codons for drug-induced UPR targets (Travers et al. 2000) in either early or late pre-meiotic cells (grown in BYTA), plotted is the average of two replicates. Hac1 signal is normalized by CPM. At bottom are traces of each gene, with colorimetric scale indicating amount of Hac1 binding, with yellow as high and blue as low. (I) Similar analysis as in (H) but showing Hac1 ChIP signal over all CDSes, plotted is the average of two replicates. Hac1 signal is normalized by CPM. (J) At top are metagene analysis of Hac1 binding around CDS start codons for targets highlighted in green in Figure 5C in either early or late pre-meiotic cells (grown in BYTA), at bottom are traces of each gene, with colorimetric scale indicating amount of Hac1 binding, with yellow as high and blue as low. (K) At top are metagene analysis of Hac1 binding around CDS start codons for targets highlighted in orange in Figure 5C in either early or late pre-meiotic cells (grown in BYTA), at bottom are traces of each gene, with colorimetric scale indicating amount of Hac1 binding, with yellow as high and blue as low. (L) Hac1 ChIP signal around the *KAR2* and *ERO1* loci with two replicates plotted (r1 and r2).

**Figure S3:**
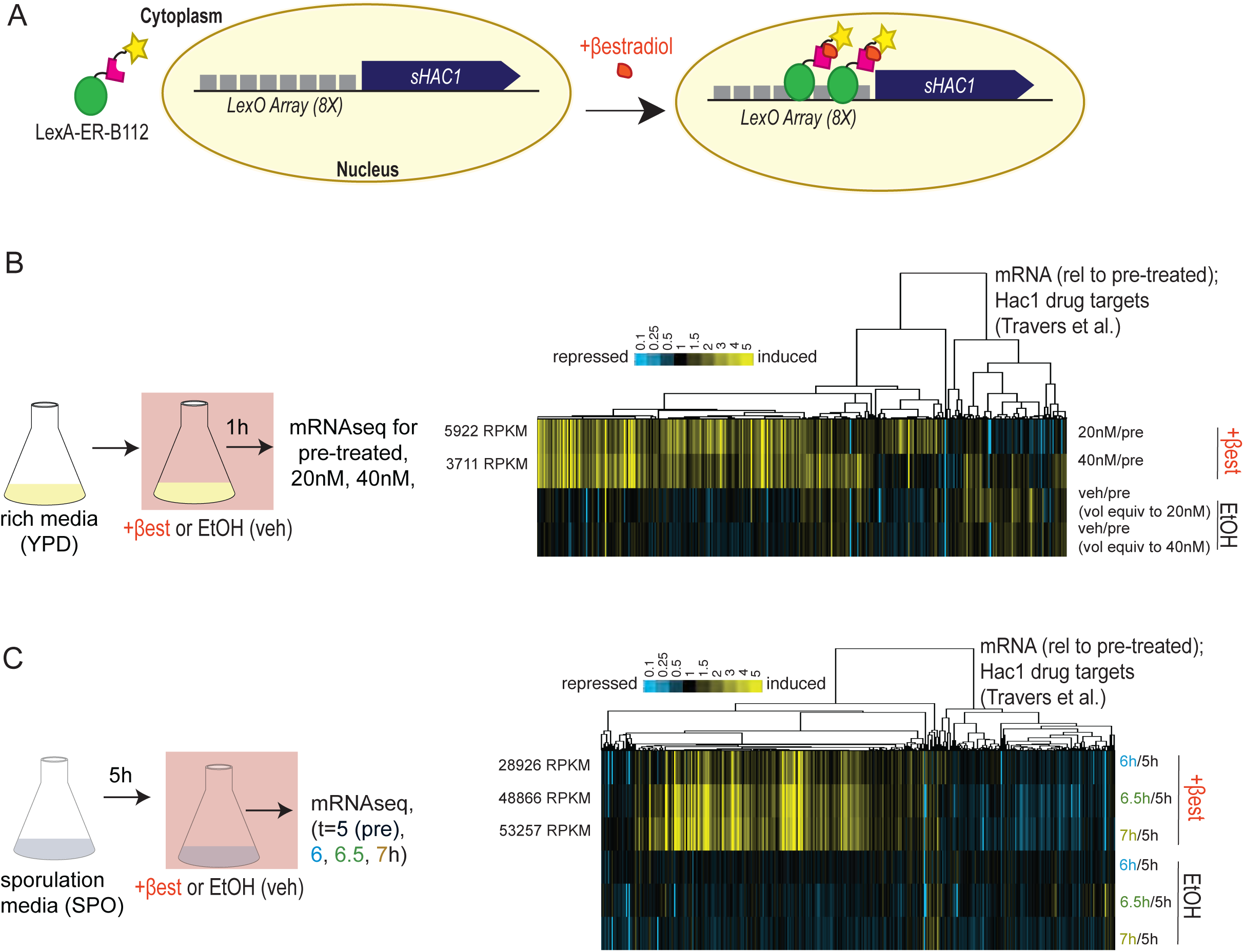
Increased *sHAC1* expression drives expression of majority of drug-induced UPR transcriptional targets. (A) Cartoon scheme for *sHAC1* overexpression approach. (B) Cartoon scheme of the mitotic experiment depicted to the right with mRNAseq data subjected to hierarchical clustering and normalized across all timepoints. RPKM for *HAC1* are shown on the left for induced samples. (C) Cartoon scheme of the meiotic experiment depicted to the right with mRNAseq data Subjected to hierarchical clustering and normalized across all timepoints. RPKM for *HAC1* are shown on the left for induced samples.

**Figure S4:**
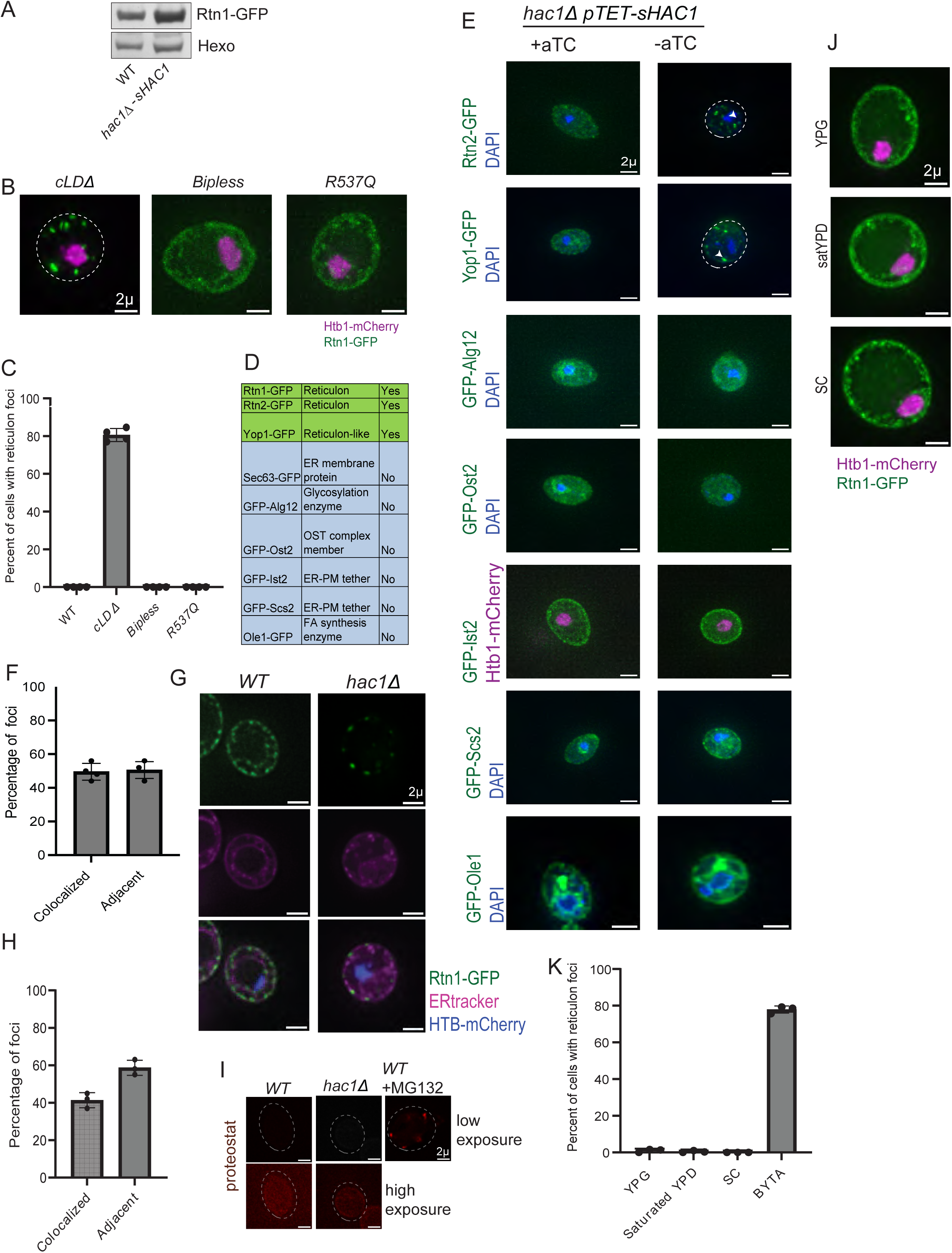
UPR-deficient puncta are specific to reticulon proteins in pre-meiotic conditions (A) Representative western blot that was quantified in Figure 4B. Hexo is Hexokinase and Rtn1-GFP is probed against anti GFP antibody. (B) Imaging of Rtn1-GFP and Htb1-mCherry in cells carrying various *ire1* allele in pre-meiotic conditions. (C) Quantification of imaging as in (B). 300 cells counted per condition per experiment, n=4. (D) Summary of presence or absence puncta when imaging various GFP-tagged ER proteins in *hac1Δ pTET-sHAC1* cells without aTC in pre-meiotic media. (E) Imaging of various ER markers in *hac1Δ* cells carrying *pTET-sHAC1* and either treated or not treated with 250ng/mL aTC for 18 hours. Strain imaged for Ist2 has Htb1-mCherry signal for labeling histones otherwise DAPI was used to stain nuclei. (F) Quantification of images as shown in Figure 4D. 300 puncta counted per condition per experiment, n=3. (G) Pre-meiotic cells were imaged as in (E), after 30 minutes of incubation at 30 degrees with 1uM ER-tracker Blue-Whtie DPX dye. (H) Quantification of images as in (G). 100 foci were counted per condition per experiment, n=3. (I) Pre-meiotic cells were imaged following fixation and subsequent incubation with proteostat reagent for 30 minutes. Treatment of WT cells with 50uM MG132 for 1 hour prior to fixation was used as a positive control. Contrast was matched in the first row but was far exceeded in the second row to show absence of any puncta structures. (J) Imaging of *hac1Δ* cells in various nutritional media. (K) Quantification of imaging as in (K). 300 cells counted per condition per experiment, n=3.

**Figure S5:**
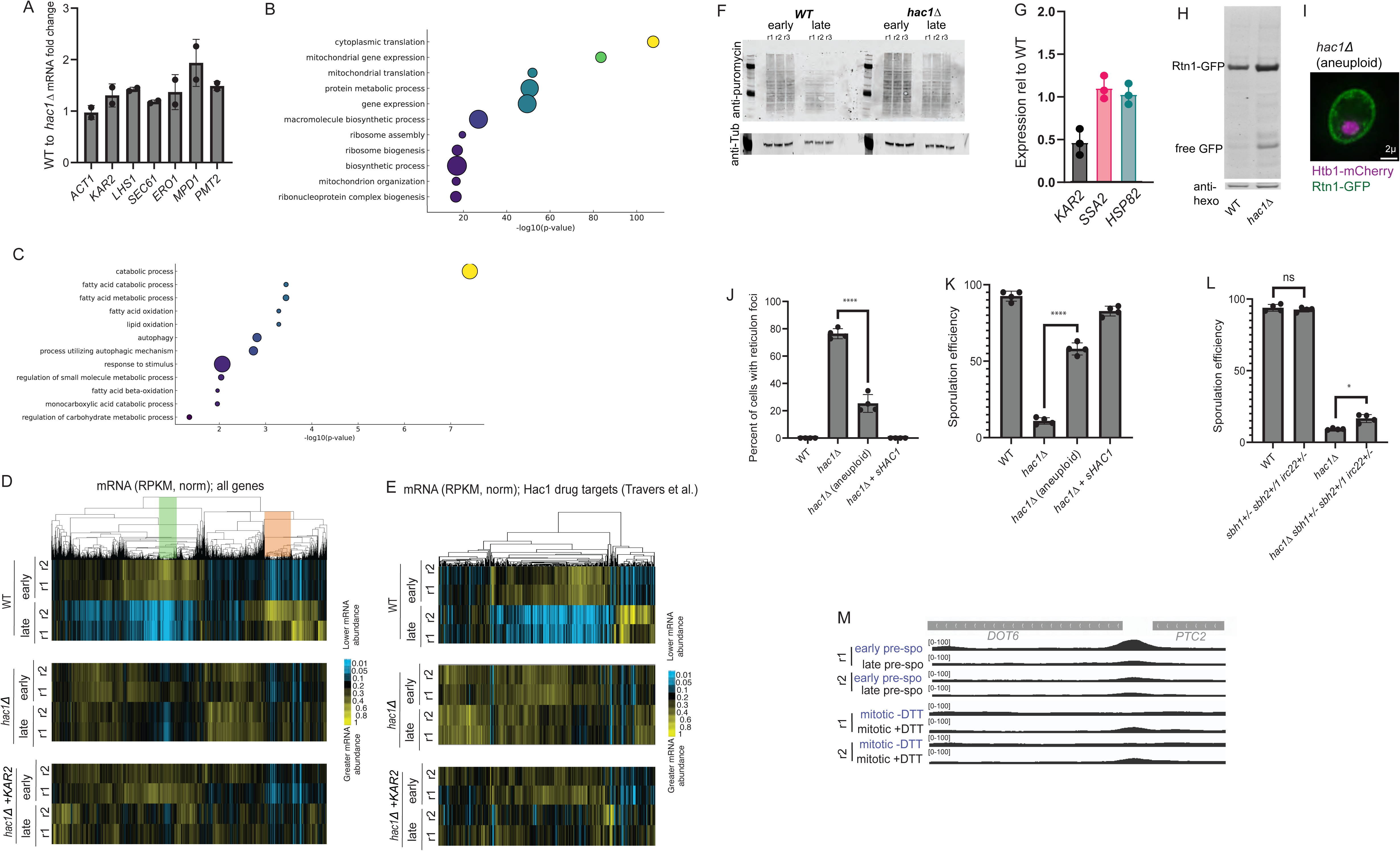
The UPR output in BYTA and meiosis is non-canonical and boosting ER folding capacity increases meiotic entry for UPR-deficient cells (A) Ratio of mRNA (RPKM) for ACT1 (control) and several known drug-induced UPR targets that are induced in early BYTA in WT cells relative to *hac1Δ*. Note mild inductions, even for the small number of targets that show effects. (B) Gene Ontology (GO) term analysis for biological processes for genes indicated by the green box in Figure 5C. (C) GO Term analysis for cellular process for transcripts in the orange box in Figure 5C (induced late in pre-meiotic UPR). (D) mRNA seq data from the experiment in Figure 5A, including *hac1Δ* cells with Kar2 expression (with *pTET-KAR2* +aTC). All data were subjected to hierarchical clustering and normalized across all timepoints. The green and orange box indicate highly differentially expressed transcript patterns in the presence and absence of Hac1 from Figure 5E. (E) mRNA seq data from the experiment in Figure 5A, including *hac1Δ* cells with Kar2 expression (with *pTET-KAR2* +aTC). Data for drug-induced UPR targets were subjected to hierarchical clustering and normalized across all timepoints. (F) Blots of puromycin incorporation, as quantified in Figure 6E. 3 biological replicates are depicted per condition and strain (G) qPCR for known Hsf1 target transcripts (including *KAR2*) in *ZPR1+/-* cells. n=3 qPCR technical replicates shown. Target expression was normalized to actin prior to fold change calculated relative to WT. (H) Representative blot of free GFP and Rtn1-GFP in pre-meiotic cells, as quantified for replicates in Figure 7G. Note that blot shown and data analyzed are same as in Figure 4B, S4A. (I) Imaging of aneuploid *hac1Δ* cells with 1 copy of chromosome V missing. (J) Quantification of imaging as in (C). 300 cells counted per condition per experiment, n=4. (K) Sporulation efficiency as scored by microscopy following 24 hours in sporulation media. 300 cells counted per strain/condition per experiment, minimum of n=4. (L) Sporulation efficiency as scored by microscopy following 24 hours in sporulation media. 300 cells counted per strain/condition per experiment, minimum of n=4. (M) Hac1 ChIP signal around the *DOT6* locus. Data from two biological repliactes are shown. Significance was determined from unpaired t-test with Welch’s correction. P-value significance is **** < 0.0001, *** < 0.001, ** < 0.01., * < .05.

## Supplementary files

File S1: Global mRNA-seq data for strains lacking *HAC1* during meiosis, pre-meiotic starvation, mitosis, and mitosis with folding inhibition. mRNAseq data quantified for two replicates for all timepoints. Data shown is recorded as RPKM (Reads Per Kilobase Million).

File S2: Global ChIP-seq data for Hac1 during meiosis, pre-meiotic starvation, mitosis, and mitosis with folding inhibition. ChIPs-eq data quantified for two replicates for all timepoints. Data shown is –log_10_qvalue of ChIP fold change with background enrichment (see methods). Nearby promoters are included along with genomic position information for each peak.

File S3: Global mRNA-seq data for strains with and without -overexpressed *sHAC1* during meiosis. Data shown is recorded as RPKM (Reads Per Kilobase Million).

File S4: Global mRNA-seq data for strains with and without over-expressed *sHAC1* during unstressed mitotic growth. Data shown is recorded as RPKM (Reads Per Kilobase Million).

File S5: Label free mass spectrometry data for early (exponential, 6h) and late (saturated, 21h) pre-meiotic growth. Normalized intensity values are shown.

File S6: Strains, plasmids, and primers used in this study. All strains used in this study are organized by strain number, strain background, and genotype. All plasmids used in this study for strain generation or experiment purposes are organized by name and description. All primers used in this study for strain or plasmid generation are organized by description and sequence.

## Materials and Methods

### Yeast strains and plasmids

All strain genotypes and plasmids used in this study are listed in File S6. The following alleles used for the microscopy experiments were first used in previous studies: Htb1-mCherry (Matos et al., 2008); Rtn1-GFP, Sec63-GFP, GFP-Scs2, and GFP-Ist2 (Otto et al., 2021). All cassette integration-based manipulations (see below) were done via restriction enzyme cutting of single integration plasmids with PmeI. Any strains or plasmids are available on request.

### Media and growth conditions

All yeast strains were grown in YPD (1% yeast extract, 2% peptone, 2% glucose) for mitotic liquid experiments, unless stated otherwise. Strains were grown on YPD plates plus 250ng/ml of aTC for maintaining euploid state. Pre-meiotic starvation was done with nonfermentable carbon source growth media (BYTA, 1% yeast extract, 2% bacto tryptone, 1% potassium acetate, and 50mM potassium phthalate) as the liquid media unless otherwise stated, with .25OD600 used as the starting inoculation concentration from saturated mitotic liquid cultures. For meiotic experiments, saturated BYTA growth cultures were pelleted, washed with sterile MilliQ water and resuspended to OD600 = 1.85 in sporulation medium (SPO; 2% potassium acetate, 40 mg/L adenine, 40 mg/L uracil, 10 mg/L histidine, 10 mg/L leucine and 10 mg/L tryptophan adjusted to pH 7.0). Cultures were allowed to shake at 30° C for the duration of the experiment. For each stage, culture volume was 1/10th of the flask volume to ensure proper aeration.

Meiotic staging:

Meiotic staging was performed scoring DAPI and tubulin morphology by fluorescent microscopy as described in Sawyer et. al. 2019. Samples were fixed in 3.7% formaldehyde for 24 hr at 4°C. Cells were then washed with 100 mM potassium phosphate pH6.4, once with sorbitol citrate (100 mM potassium phosphate pH7.5, 1.2 M sorbitol), and digested in 200 μL sorbitol citrate, 20 μL glusulase (Perkin-Elmer) and 6 μL of zymolase (10 mg/mL, MP Biomedicals) for 3 hours at 30° C while rotating. Samples were pelleted at 900 rcf for 2 min, washed with 100 μL sorbitol citrate, pelleted again and resuspended in 50 μL sorbitol citrate. Samples were then mounted on slides prepared with poly-L-lysine, submerged in 100% methanol at -20° C for 3 min, transferred to 100% acetone at -20° C for 10 sec, then allowed to air dry. Samples were then incubated at RT for 1 hr in primary anti-tubulin antibody (Bio-Rad, 1:200) in PBS-BSA (5 mM potassium phosphate, 15 mM NaCl, 1% BSA, 0.1% sodium azide). Samples were then washed 3x in PBS-BSA and incubated with preabsorbed FITC-conjugated secondary antibody (Jackson ImmunoResearch Labs, 6:200) for 1 hr at RT. Samples were washed 3x with PBS-BSA and mounted with VectaShield Antifade Mounting Medium with DAPI (Vector Labs).

### Strain generation

Standard PCR based homologous recombination cassette swapping strategies in yeast were used to generate deletions of genes for this study (Longtine et al., 1998; Powers et al., 2022). Cas9-mediated deletion of the lumenal domain of Ire1 was done by co-transforming cells with a selectable Cas9 expression plasmid and a double-stranded repair template. To generate euploid UPR-deficient strains, deletion of diploids as well as haploids was performed via transformation of selection cassettes into strains carrying genomically integrated aTC-inducible alleles of either *HAC1, ROT1,* or *KAR2* integrated. Strains were plated post-transformation on plates with aTC such that all strains were constantly in the presence of UPR activity mimetics. Heterozygous diploids were then dissected to generate haploids to backcross for construction of stable haploid or diploid lines. Importantly, strains were always grown on aTC plates except for when experiments required a no UPR condition.

### Growth Curve Analysis

Plates were sealed with a Breathe Easy Cover (Sigma-Aldrich) and grown at 30C for 24 hours in a plate reader. Absorbance readings were collected every 15 minutes by absorbance at 600 nm, with agitation before each reading. All experiments were loaded in triplicate in the 96-well plate.

### Fluorescent Microscopy

Images were acquired using a DeltaVision Elite wide-field fluorescence microscope (GEHealthcare, Sunnyvale, CA). Timelapse microscopy was performed exactly as described in King et al., 2019. When timelapse was done with BYTA instead of SPO, similar to with conditioned SPO being used instead of fresh SPO, conditioned BYTA was used instead of fresh BYTA. All time-lapse experiments were performed using the CellAsic system (EMD Millipore) in Y04D or Y04E microfluidics plates. Images were deconvolved using softWoRx imaging software (GE Life Sciences).

For linescan analysis of microscopy a 1 micron line was drawn by eye over the middle where a clear Rtn1-mCherry foci was localized in FIJI image analysis software. The surface plot tool was used to quantify the fluorescent intensity for both channels and this was plotted along the length of the line.

The following microscopy settings were used for the respective alleles:

Sec63-GFP was imaged using the following settings:100%T, .2s EX: EX: 475/28 EM: 523/36

All other GFP alleles were imaged using: 32%T, 0.025s EX: 475/28 EM: 523/36.

Htb1-mCherry was imaged using: 10%T, 0.025s EX:575/25 EM: 632/60.

Rtn1-mCherry was imaged using: 32%T, 0.025s EX:575/25 EM: 632/60.

ERtracker was imaged using: 100%T, 0.1s EX:390/18 EM: 435/48.

Proteostat was imaged using: 32%T, 0.1s EX:575/25 EM: 632/60.

DAPI was imaged using: 100% T, .1s EX: 390/18 EM: 435/48

For timelapse microscopy the movies were taken with 8 microns of z across 8 steps while the non-timelapse microscopy was done with 2.5 microns of z across 12 steps.

### Proteostat assay staining

Proteostat reagent staining was done as is reported in Gestaut et al., 2019 aside from a couple small minor differences. Cells in this study were grown at 30 degrees in BYTA for 21 hours before fixation. After proteostat addition the cells were incubated at room temperature instead of 4 degrees for 30 minutes. Cells were imaged with the above settings.

### RNA extraction

Cells were harvested via pelleting liquid culture at 6000 RCF then aspirating media before snap freezing in liquid nitrogen. Cells were thawed on ice and resuspended in 625ul TES buffer (10 mM Tris pH 7.5, 10 mM EDTA, 0.5% SDS). An equal volume of citrate-buffered acid phenol (pH 4.3, P4682, Sigma-Aldrich) was added to cells, and they were incubated at 65C for 1 hour in a Thermomixer C (Eppendorf) shaking at 1400 RPM. After microcentrifugation (20,000 g for 10 min) the aqueous phase was transferred to a second tube with 600ul acid phenol. The aqueous phase was separated by microcentrifugation (20,000 g for 10 min) and transferred to a third tube with 600ul chloroform. The aqueous phase was separated by microcentrifugation (20,000 g for 10 min) and RNA was precipitated in 100% isopropanol with 350 mM sodium acetate (pH 5.2) overnight at –20C. Pellets were washed with 80% ethanol and resuspended in DEPC water.

### RT-qPCR

cDNA synthesis was done exactly as detailed in Morse et al., 2024. Only one difference was done compared to that protocol: the quantification was performed using 4.8ul of 1:16 diluted cDNA instead of 1:20 diluted cDNA.

### Northern blotting

All northern blotting was followed exactly as described in Van Dalfsen et al., 2018 except for a few minor differences. The agarose gel was run for 3 hours at 100V. The low stringency washes were done for 15 minutes each with 3 washes. The high stringency washes were done for 15 minutes each with 3 washes.

### mRNA sequencing

Libraries were constructed as in Morse et al., 2024. RPKM (Reads per kilobase million) were calculated for all samples.

### Immunoblotting

Cell lysis, protein isolation, and immunoblotting protocol was performed as reported in Morse et al., 2024. Primary antibodies: mouse anti v5 (R960-25, *Thermo Fischer Scientific*) used at a concentration of 1:2000, rabbit anti Kar2 (gift from Mark Rose) used at a concentration of 1:2000, rabbit anti Hexokinase (H2035, *US Biological*) used at a concentration of 1:5000, rat anti Tubulin (Serotec, 1:10,000). Secondary antibodies: donkey anti mouse antibody conjugated to IRDye 800CW used at a concentration of 1:15,000 (LI-COR); goat anti rabbit secondary conjugated to IRDye 680RD at a concentration of 1:15,000 (LI-COR), goat anti rat secondary conjugated to IRDye 680RD at a concentration of 1:15,000 (LICOR).

### Puromycin incorporation

Cells were treated with 250uM puromycin for 30 minutes before protein lysis was performed as in Morse et al., 2024. Mouse anti puromycin (Millipore) was used at a concentration of 1:25,000.

### Mass spectrometry sample prep

A total of 40 μg of protein from cell lysates, quantified using the Pierce™ BCA Protein Assay Kit (Thermo Scientific) according to the manufacturer’s instructions, was reduced with 5 mM dithiothreitol (DTT) for 45 minutes at 25 °C with shaking at 600 rpm. Reduction was followed by alkylation with 10 mM iodoacetamide (IAA) for 45 minutes in the dark under the same conditions. Samples were then diluted with 50 mM Tris-HCl (pH 8.0; Thermo Scientific) to reduce the urea concentration to below 2 M. Proteins were digested overnight at 25 °C and 600 rpm using 0.8 μg of sequencing-grade modified trypsin (Promega). After digestion, peptides were acidified with formic acid (Thermo Scientific) and desalted using in-house packed C18 StageTips (two plugs), following the protocol described by Rappsilber et al, 2007. Cleaned peptides were dried using a Thermo Savant SpeedVac and reconstituted in 3% acetonitrile/0.2% formic acid prior to LC-MS/MS analysis.

### LC-MS/MS analysis on a Q-Exactive HF

Approximately 1 μg of total peptides were analyzed on a Waters M-Class UPLC using a 15 cm x 75 µm IonOpticks C18 1.7 µm column coupled to a benchtop Thermo Fisher Scientific Orbitrap Q Exactive HF mass spectrometer. Peptides were separated at a 400 nL/min flow rate with a 90-minute gradient, including sample loading and column equilibration times, using solvents A (0.1% formic acid in water) and B (0.1% formic acid in acetonitrile). The detailed gradients of solvent B are below: 2% B for 1 min; linear increase to 10% B over 29 min; linear increase to 22% B over 27 min; linear increase to 30% B over 5 min; linear increase to 60% B over 4 min; linear increase to 90% B over 1 min, held for 2 min; linear decrease to 50% B over 1 min, held for 5 min; linear decrease to 2% B over 1 min; and re-equilibrated at 2% B for 14 min.

Data were acquired in data-independent mode using Xcalibur software (4.5.474.0). MS1 spectra were measured with a resolution of 120,000, an AGC target of 3e6, and a scan range from 350 to 1600 m/z. 21 isolation windows of 60 m/z were measured at a resolution of 30,000, an AGC target of 3e6, normalized collision energies of 22.5, 25, 27.5, and a fixed first mass of 200 m/z.

Raw DIA data were processed using DIA-NN (version 2.1.0) (Demichev et al, 2020). DIA-NN was used in library-free mode, in which an in silico spectral library was generated directly from the *Saccharomyces cerevisiae* protein sequence database (UniProt proteome: UP000002311). The predicted spectral library was then used for peptide identification and protein quantification across all samples. The analysis were performed using default DIA-NN settings, with protein N-terminal acetylation as variable modification, match between runs enabled and protein inference enabled.

### Chromatin Immunoprecipitation

For the vegetative ChIP collection, a YPD culture was started at an OD600 of 0.2 and grown to an OD600 of approximately 0.4 at 30°C, split into two flasks, and 4.5mM DTT was added to one flask. After 30 minutes of additional growth, approximately 90 OD600 units of cells were harvested for ChIP. Cells were fixed using fresh 1% formaldehyde (Sigma Aldrich 252549) with periodic inversion at room temperature for 15 minutes. The crosslinking reactions were quenched using 125mM glycine (Fisher BP381-1) with light shaking for 5 minutes. Cells were pelleted at 4°C, then washed using ice cold PBS and pelleted again. Supernatant was discarded and cell pellets were resuspended in cold FA lysis buffer (50 mM Hepes pH 7.5 (Fisher BP310-1), 150 mM NaCl, 1 mM EDTA, 1% Triton (Sigma Aldrich T9284), 0.1% sodium deoxycholate (Sigma Aldrich D6750)) with 0.1% SDS and protease inhibitors (Roche cOmplete protease inhibitor cocktail, PMSF, Pepstatin A). SDS was added fresh to every buffer from a 20% stock solution. Samples were centrifuged, supernatant was discarded, and pellets were flash frozen in liquid nitrogen. For the BYTA ChIP collection, a BYTA culture was started at an OD600 of 0.2 and grown at 30°C. Approximately 90 OD600 units of cells were harvested at 6 hour and 21 hour timepoints. Cells were treated as above. For the meiotic ChIP collection, SPO cultures were set up at on OD600of 1.9 at 30°C. Approximately 170 OD600 units of cells were harvested at 5 hour 30 minutes, 6 hour 45 minutes, and 8 hour timepoints. Cells were treated as above.

Approximately 500uL Zirconia beads (Biospec 11079105z) were added to cell pellets, and cells were resuspended in cold FA lysis buffer with 0.1% SDS and protease inhibitors and lysed by Beadbeater (Mini-Beadbeater-96, Biospec Products). Lysates were collected by centrifuging at 500xg for 1 min. All centrifugation steps were carried out at 4°C. Cell debris was pelleted at 2000xg for 3 min. Supernatants were retrieved and centrifuged at 20,000xg for 15 minutes to pellet chromatin. Chromatin pellets were resuspended in cold FA lysis buffer with 0.1% SDS and protease inhibitors and moved to 15 mL sonication tubes containing 300 uL sonication beads (Diagenode C01020031). Samples were sonicated in a Bioruptor Pico (Diagenode) for 30 seconds on / 30 seconds off for 8 cycles to generate fragments around 150–400 bp. Samples were centrifuged at 20,000xg for 1 min and supernatant was saved for use in the immunoprecipitation (IP).

For the IP, 40 uL mouse anti-3V5 agarose slurry (Millipore Sigma A7345) was washed 3 times using cold FA lysis buffer with 0.1% SDS. For each wash, agarose beads were nutated for 5 min at 4°C, then pelleted by centrifugation at 1,000xg for 30 seconds. 30 uL of sheared chromatin was set aside prior to the IP as input. The remaining chromatin was added to the washed anti-3V5 beads and nutated overnight at 4°C. The next day, beads were washed twice with FA lysis buffer with 0.1% SDS, twice with a high salt buffer (FA lysis buffer with 0.1% SDS, and 250 mM NaCl), and twice with a high detergent buffer (10 mM Tris pH 8, 250 mM LiCl, 0.5% NP-40 (Millipore Sigma 56741), 0.5% sodium deoxycholate, and 1 mM EDTA). 130uL of TE (10 mM Tris pH 8, 1 mM EDTA) with 1% SDS was added to both the IPs and inputs. Samples were eluted by shaking in a thermomixer (Eppendorf) at 450 RPM and 65°C overnight. Samples were cleaned up using QIAQuick PCR Purification Kit (Qiagen 28104).

### ChIP sequencing

Sequencing libraries were generated using the NEXTFLEX Rapid DNA-Seq kit 2.0 according to manufacturer’s instructions (Revvity NOVA 5188-02). SPRIselect beads were used to select for fragments between 200-700 bp (Beckman Coulter B23318). Libraries were quantified using a Qubit fluorometer (Invitrogen), and library size was checked using the Agilent 4200 TapeStation (Agilent Technologies, Inc). Samples were submitted for 150 bp paired-end sequencing on the NovaSeq X at the Vincent J. Coates Genomics Sequencing Laboratory (QB3 Genomics, UC Berkeley, Berkeley, CA, RRID:SCR_022170).

### ChIPseq analysis

Reads from input and IP samples were aligned to the SK1 genome using Bowtie2 (version 2.3.4.1; Langmead and Salzberg, 2012) and peaks were called using MACS2 (version 2.2.9.1; Zhang et al., 2008). Peaks were annotated using ChIPseeker (version 1.44.0; Yu et al., 2015) if the peak summit overlapped with a promoter (defined here as 600 bp upstream of gene). Only peaks meeting a fold enrichment (ChIP over input) cut-off of ≥ 4 were used in the analysis. Normalized bigWig files were generated for visualization in IGV and metagene analysis using deepTools (version 3.5.6) bamCoverage and replicates were averaged using bigwigCompare. Metagene analysis was performed using deepTools computeMatrix reference-point and visualized using plotHeatmap and plotProfile (Ramírez et al., 2016).

## Data accessibility

All raw mRNAseq and ChIPseq data will be made available publicly at the time of publication.

